# A Multi-Objective Scoring (MOS) Framework for Detecting Cross-Modal Spatial Similarity: Conceptual and Direct Formulations

**DOI:** 10.64898/2026.04.09.717471

**Authors:** Amin Jarrahi, Allison R. Jones, Weisheng Tang, Hairong Qi, A. Colleen Crouch

## Abstract

Spatial multi-omics methods allow researchers to study complex biological systems by integrating multiple molecular layers while preserving their spatial organization. However, integrating spatial transcriptomics and mass spectrometry imaging (MSI) remains challenging due to the differences between the two modalities, including sampling geometry, spatial resolution, signal scaling, and measurement principles. For example, 10x Genomics Visium captures transcriptomic data on a discrete hexagonal grid of spots; however, matrix-assisted laser desorption/ionization mass spectrometry imaging (MALDI-MSI) produces dense Cartesian pixel maps of molecular distributions. These differences make exact spatial co-registration difficult and limit the effectiveness of single-metric similarity approaches.

Here, we introduce a Multi-Objective Scoring (MOS) framework designed to detect cross-modal spatial similarity without requiring exact pixel-to-pixel alignment. The MOS framework integrates multiple complementary spatial descriptors into a unified similarity score, including coordinate-based metrics (value correlation, importance-based Intersection-over-Union (IoU), and importance-map correlation) and descriptor-based metrics that capture higher-order spatial organization, such as spatial histograms, radial profiles, quadrant statistics, and Moran’s I spatial autocorrelation. These metrics are combined through a weighted ensemble model. This model calculates the weights using synthetic spatial datasets that simulate realistic tissue geometry, sampling differences, and spatial distortions.

The framework was applied to a spatial multi-omics dataset from murine brain tissue, integrating spatial transcriptomics with MALDI-MSI lipidomics across young and aged control and Alzheimer’s disease (AD) models. Synthetic data validation results demonstrated strong pattern-matching performance (96.14% accuracy), and application to experimental data identified several MSI analyte features whose spatial distributions closely matched transcriptomic patterns. In particular, strong and reproducible associations were observed between myelin-related genes (*Mbp* and *Plp1*) and multiple analyte features enriched in white matter regions.

Overall, whether applied conceptually or directly, the MOS framework provides a strong strategy for cross-modal spatial integration and offers a scalable tool for discovering spatial relationships across diverse multi-omics datasets and facilitating hypothesis generation.

## Introduction

Spatial multi-omics has become a powerful approach for exploring complex biological systems by integrating corresponding molecular layers. This can be achieved either by profiling multiple modalities on the same tissue section or by aligning consecutive, closely matched sections to maintain spatial continuity [1], [2], [3], [4], [5]. With platforms such as 10x Genomics Visium Spatial Gene Expression, researchers can characterize gene expression patterns of tissue through spatial transcriptomics[6], [7]. Similarly, mass spectrometry imaging (MSI), including matrix-assisted laser desorption/ionization (MALDI-MSI), enables spatially resolved mapping of other biomolecules, including lipids, directly from tissue sections[8], [9], [10], [11], [12], [13]. Combining these technologies together provides a more extensive and complementary molecular view of tissue structure at high spatial resolution[4], [5].

Computational approaches for multi-omics integration have evolved from non-spatial models to frameworks that incorporate tissue architecture [5]. Early integrative methods, such as Multi-Omics Factor Analysis (MOFA) and its successor MOFA+, established foundational approaches for identifying major patterns across molecular layers by understanding underlying sources of variation [14], [15]. These models effectively separate shared and modality-specific sources of heterogeneity. However, these models were primarily developed for non-spatial single-cell data and do not explicitly incorporate anatomical coordinates. To address this limitation, more recent methods such as COSMOS in 2025 and MAGPIE in 2026 have been introduced. COSMOS uses graph neural networks to integrate paired spatial datasets. It can capture complementary features while preserving shared spatial coordinates [16]. Similarly, the MAGPIE framework was developed to support the co-registration of spatial transcriptomics and metabolomics data from either the same or consecutive tissue sections. This framework emphasizes the importance of mapping different modalities into a shared coordinate space to enable cross-modal analysis [4].

With significant progress in spatial profiling technologies, there are still major methodological challenges in robust integration of spatial transcriptomics and MALDI-MSI datasets [16]. There are significant differences in these modalities including sampling geometry, spatial resolution, signal scaling, noise structure, and underlying physical measurement principles. Among these differences, the mismatch in grid architecture poses one of the most substantial obstacles and further intensifies resolution disparities. The 10x Genomics Visium platform employs a hexagonal array of discrete capture spots with defined center-to-center spacing for the Visium Gene Expression Slides. These spots do not form a continuous sampling surface, and gaps exist between neighboring capture locations. Therefore, it creates a partially discrete spatial representation of the tissue. The overall coverage of spots is less than 30% of the capturing area; however, the use of the hexagonal grid is more efficient compared to rectangular grid by reducing the sampling size by 13.4% [17], [18]. In contrast, MALDI-MSI produces data on a dense Cartesian grid in which adjacent pixels are tightly connected. Therefore, exact spatial alignment between modalities is rarely achievable in practice, and relying on a single similarity metric fails to adequately represent the complex, multidimensional structure of biological spatial relationships[4].

The importance of robust integration becomes especially evident when considering complex molecular interactions that drive many diseases. Spatial multi-omics is particularly valuable for revealing how intracellular processes are formed by, and interact with, the surrounding microenvironment. These processes become more important in areas such as tumor immunology, developmental biology, and tissue injury. For example, integrated analyses have linked metabolic changes – such as increased lactate production – to specific transcriptomic signatures in damage-associated macrophages during pulmonary fibrosis [4]. In oncology, mapping gene expression alongside metabolic gradients has revealed how changes in the extracellular matrix and alterations in cellular metabolism work together to drive tumor progression and enable immune evasion [4]. Considering these advancements in this field, major challenges remain unsolved due to fundamental differences in sampling geometry and measurement principles between technologies such as spatial transcriptomics and mass spectrometry. This highlights the need for generalized, geometry-aware frameworks that can quantify spatial similarity without relying on perfect co-registration across modalities.

We present a Multi-Objective Scoring (MOS) Framework designed to quantify spatial similarity between heterogeneous omics modalities without requiring perfect co-registration. This framework is designed to remain robust in the presence of minor spatial misalignments and orientation differences, while adapting to variations in data quality. It integrates complementary geometric and statistical descriptors, including radial profiles, spatial histograms, quadrant statistics, spatial autocorrelation (Moran’s I), and intensity-distribution similarity, to capture both direct spatial correspondence and higher-order structural relationships. The component metrics are combined via a weighted ensemble model, with weights calibrated using synthetic data that systematically simulate spatial misalignment, orientation perturbations, resolution differences, acquisition variability, and realistic mouse half-brain anatomical geometry.

Our framework uses selected spatial patterns within the gene modality and subsequently searches for candidate lipid features that exhibit similar structural signatures. This targeted strategy substantially reduces computational cost compared to exhaustively evaluating cross-modal similarity for all gene–lipid combinations. We applied the framework to Alzheimer’s disease (AD) mouse brain tissue as a case study, given its extensive investigation over several decades and its relevance as a complex neurodegenerative disorder [19], [20], [21], [22]. This highlights the utility of the proposed framework in a biologically meaningful context. The main aim of this work presented here is not to generate AD-specific insights, but to design and validate a generalized, geometry-aware computational framework for cross-modal spatial integration. However, this framework can be applied either directly by using the platform as described here or conceptually using MOS and modifying metrics to fit a wide range of spatial omics combinations and provides a scalable foundation for future multi-omic investigations across tissues, organisms, and disease models, which reflects its modality-agnostic nature.

## Methods

### Sample Collection

Murine brains from young (10–12 weeks) and aged (52–54 weeks) female 3xTg-AD transgenic mice and age-matched controls (n = 4 per group) [23] were harvested and rapidly flash-frozen in liquid nitrogen to preserve tissue morphology and RNA integrity. The brains were cryosectioned coronally at 10 µm thickness using a Leica cryostat. Sections encompassing the cortex and hippocampus, at a location approximately 2.24 ± 0.21 mm from the bregma, were collected in series. To enable cross-modal spatial analysis, consecutive sections were mounted in an alternating manner: one section was mounted onto glass microscope slides (Fisher Scientific) for MALDI mass spectrometry imaging [24], [25], [26], and the immediately adjacent section was mounted onto 10x Genomics Visium Spatial Gene Expression slides for spatial transcriptomic profiling. This serial mounting strategy ensured that lipidomic and transcriptomic measurements were obtained from closely matched anatomical regions. Additionally, seven serial sections from a single animal were mounted on 10x Genomics Tissue Optimization (TO) slides for performing the Visium Tissue Optimization Protocol. This protocol was implemented to determine the optimal permeabilization time, allowing efficient mRNA release while preserving tissue morphology and ensuring high-quality, reproducible gene expression. After mounting, all slides were placed in plastic slide jars (the lids of which were tightly sealed to avoid moisture exposure), transferred in dry ice, and stored at −80 °C until further processing.

### MALDI-MSI Lipidomics

The washing, sublimating, scanning, and mass spectrometry imaging were performed at the Mass Spectrometry Research Center (MSRC) at Vanderbilt University. Samples, mounted on glass slides, were transferred in the plastic slide jars on dry ice to Vanderbilt. The samples on glass slides were submerged in four consecutive baths containing isotonic ammonium formate solution (150 mM for 45 seconds. Following the washing steps, the slides were gently dried under a continuous stream of nitrogen gas to minimize moisture and prevent analyte delocalization. To further ensure complete dehydration and stabilization of the tissue before subsequent analyses, the dried slides were placed in a vacuum desiccator for 20 minutes[27]. A matrix solution DMACA was applied through the sublimation method (HTX SubliMATE) with optimized parameters (Table 1) [27].

**Table 1.**
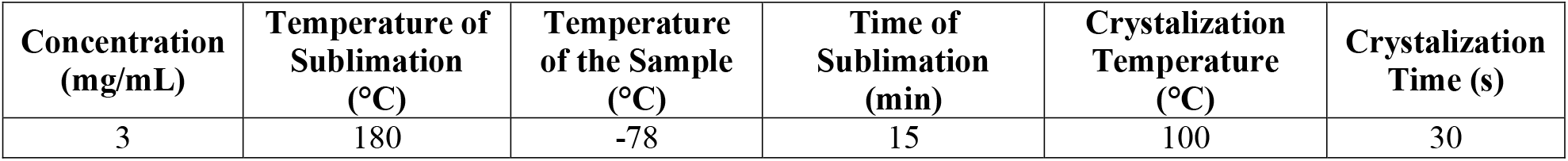
DMACA matrix sublimation conditions.

The slides were first fluorescence-scanned using a ZEISS Axio Scan.Z1 slide scanner. They were then analyzed with Bruker’s timsTOF fleX mass spectrometer under the parameters listed in Table 2. Following imaging, the samples were placed in the slide jar and vacuumed, sealed, and stored at −80 °C freezer at the University of Tennessee. Then after removing from freezer, the matrix was removed by three sequential washes in pure methanol, each lasting 1 minute. Finally, the slides were stained and imaged at 2× resolution using a Leica M205 FCA microscope at the Advanced Microscopy and Imaging Core (AMIC) of the University of Tennessee, Knoxville (Figure 1).

**Table 2.** MALDI timsTOF fleX instrumental parameters for positive ion mode.

|  |  |  |
| --- | --- | --- |
| **Transfer** | MALDI Plate Offset (V) | 50.0 |
|  | Deflection 1 Delta (V) | 70.0 |
|  | Funnel 1 RF (Vpp) | 350.0 |
|  | isCID Energy (eV) | 10.0 |
|  | Funnel 1 RF (Vpp) | 300.0 |
|  | Multipole 1 RF (Vpp) | 300.0 |
| **Collision Cell** | Collision Energy (eV) | 10.0 |
|  | Collision RF (Vpp) | 2500.0 |
| **Quadrupole** | Ion Energy (eV) | 5.0 |
|  | Low Mass (analyte) | 450.0 |
| **Focus Pre ToF** | Transfer Time ($\mu\text{s}$) | 90.0 |
| | Pre Pulse Storage ($\mu\text{s}$) | 12.0 |
| **Detection** | High Sensitivity Detection – Low Sample Amount | False |
|  | Focus Mode | False |

**Figure 1.**
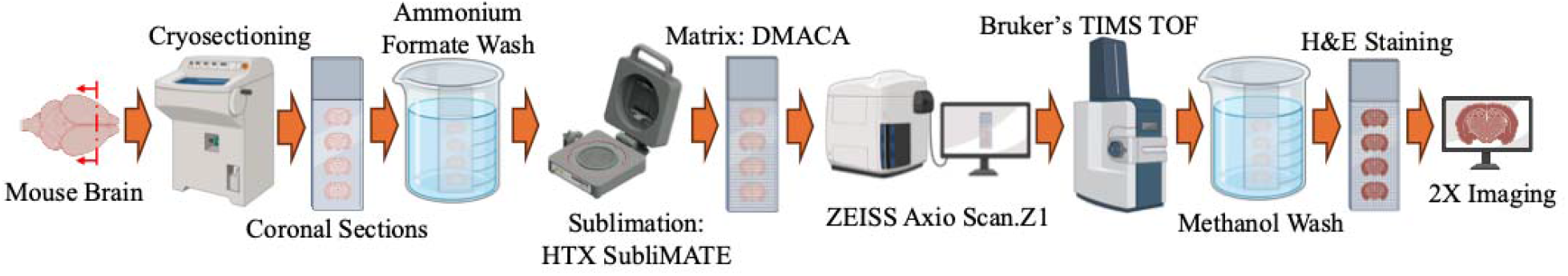
Schematic overview of the experimental workflow for MALDI-MSI, highlighting tissue preparation, matrix application, data acquisition, and imaging^a^.

For MALDI-MSI data processing, our previously published algorithm was employed and enhanced, and new features were added to the algorithm to enable it to read and analyze datasets from various companies and also to perform better peak detection and peak alignment [28]. For this study, data preprocessing involved smoothing (Savitzky–Golay filter)[29], baseline correction (asymmetric least squares, AsLS) [30], peak alignment using four lock masses [496.339 (LPC 16:0) [31], [32], 734.569 (PC 32:0) [31], [33], 760.585 (PC 34:1) [31], [32], [34], and 806.569 (PC 38:6) [34]] together with drift line correction, global peak detection (minimum peak distance = 0.024 Da), peak augmentation (± 0.012 Da or up to ± 2 indices), and total ion current (TIC) normalization. As the goal was to examine the distribution pattern of the analyte within each sample to compare across groups, only analytes detected across all sample groups were retained, and those not commonly present were excluded from further analysis. Subsequently, normalization, logarithmic transformation, Z-score scaling, principal component analysis (PCA), and k-nearest neighbors (kNN) were performed. Leiden clustering at a low resolution (res = 0.03) was then applied to identify and remove off-tissue and contaminated pixels. The processed data were stored in AnnData (.h5ad) format (Figure 2).

**Figure 2.**
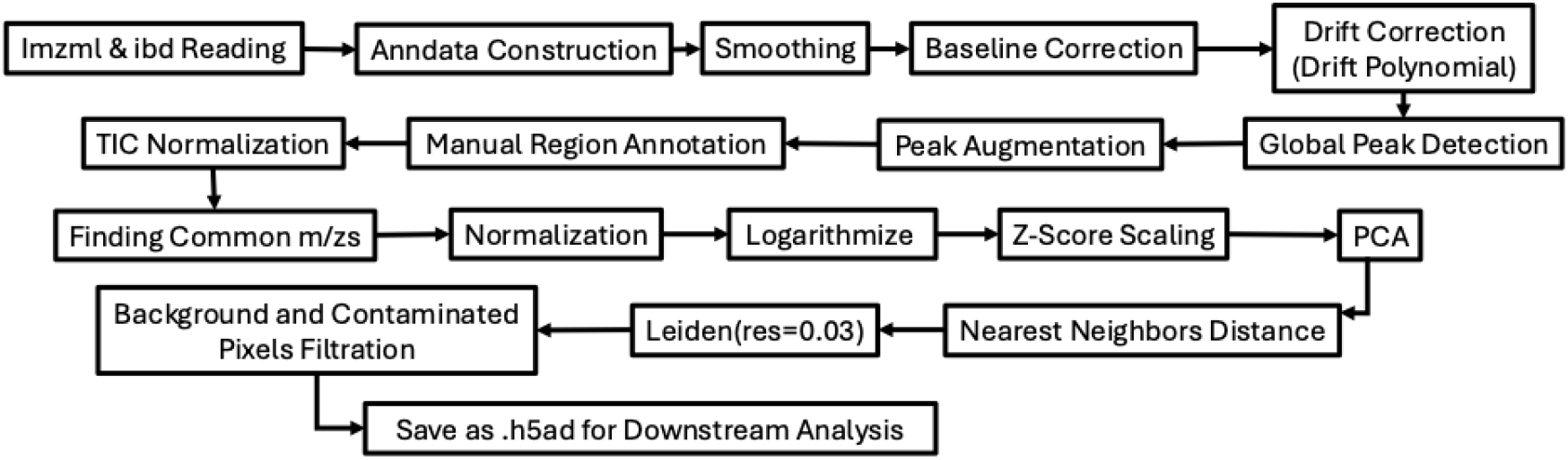
Flowchart summarizing the computational steps for MALDI-MSI data implemented in Python.

### Spatial Transcriptomics

10x Genomics Visium Spatial Gene Expression slides contained an array of spatially barcoded capture spots that enabled the assignment of transcriptomic information to precise tissue locations. The 10x Genomics Visium Spatial Gene Expression protocol was performed following the 10x Genomics Tissue Optimization (TO) protocol (Figure 3). The tissue sections were briefly warmed to ensure proper adhesion to the slide surface and were subsequently fixed to preserve tissue morphology and RNA integrity. The sections were then stained using hematoxylin and eosin to visualize tissue architecture and imaged under bright-field microscopy, generating high-resolution images to guide downstream analyses, at Advanced Microscopy and Imaging Core (AMIC) of the University of Tennessee, Knoxville. Following imaging, the slides were processed for spatial transcriptomic library preparation according to the 10x Genomics Visium protocol with a permeabilization time of 12 minutes. The resulting cDNA libraries were sent to the Genomics Core at the University of Tennessee, Knoxville, for sequencing on an Illumina platform (NovaSeq). Sequencing outputs were delivered as FASTQ files, containing raw reads indexed by spatial barcodes and ready for computational processing and spatial gene expression analysis (Figure 3).

**Figure 3.**
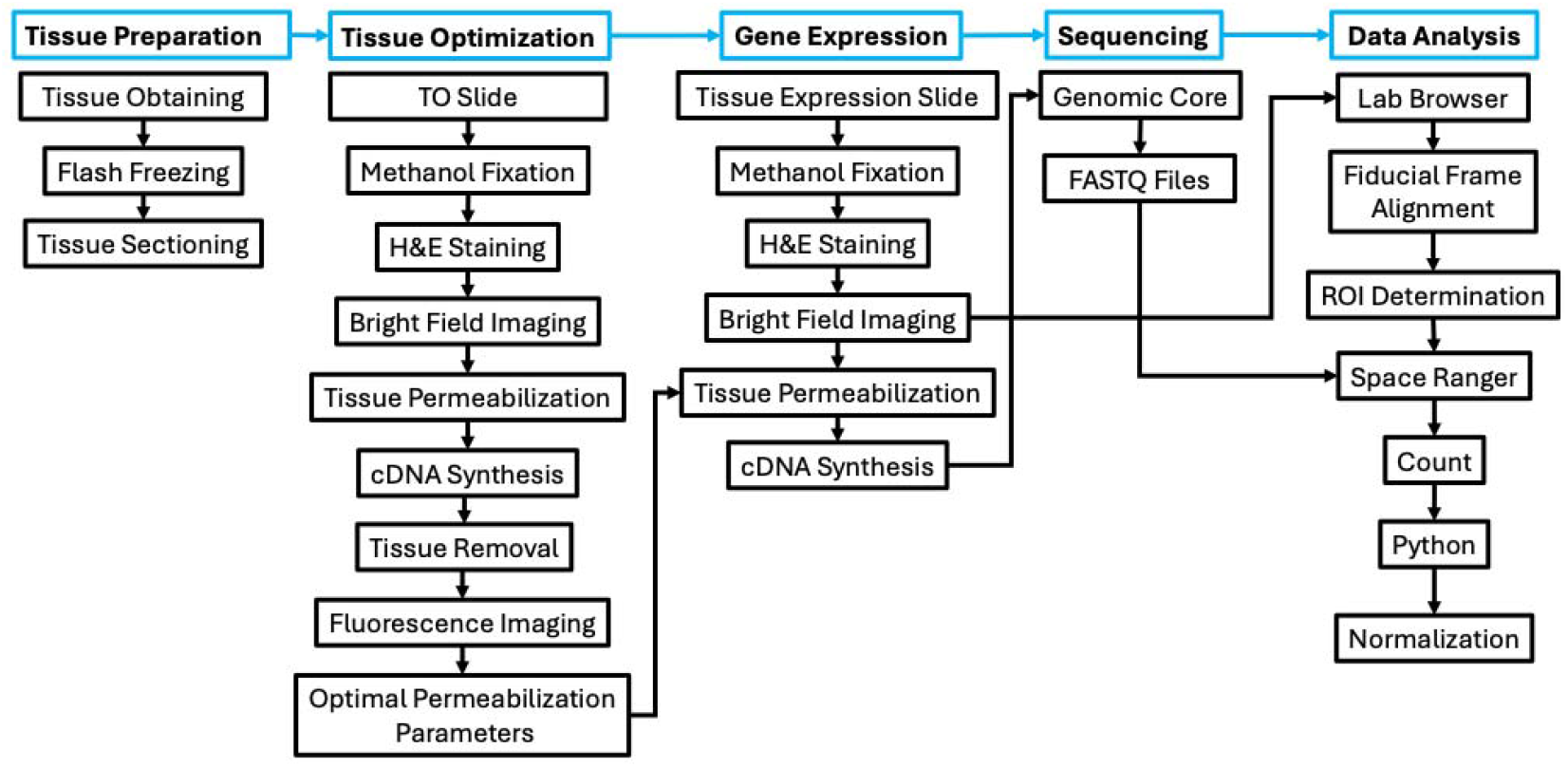
Flowchart of Visium tissue optimization, spatial gene expression analysis, and data preprocessing pipeline.*^b^* TO: tissue optimization

Brightfield H&E-stained tissue images were first examined in Loupe Browser for fiducial frame alignment and determination of the region of interest (ROI). Sequenced data were obtained as FASTQ files from the Genomic Core and were processed using 10x Genomics Space Ranger to align reads, assign them to spatial barcodes, and generate spot-level count matrices. The processed data were then imported into Python for normalization, and the spatial coordinates of each spot (X and Y, in micrometers) were incorporated into the corresponding samples’ AnnData objects (Figure 3).

No further large-scale analysis was performed on the spatial transcriptomics dataset. Instead, eight genes – *Mbp* [35], [36], *Plp1* [37], [38], *S100b* [39], *Hspa1a* [40], [41], *Mapt* [42], [43], [44], *Thy1* [45], *App* [46], [47], and *Apoe* [46], [48], [49], [50] – were selected for subsequent analysis based on previously reported evidence of their upregulation, downregulation, dysregulation, or involvement in pathological mechanisms associated with aging, AD, or both.

### Multi-Objective Scoring Framework

We employed a MOS pipeline that did not require pixel-to-pixel overlay. We focused on identifying differentially expressed genes within the spatial transcriptomic dataset and evaluated whether corresponding spatial patterns can be detected in the lipidomic images. Integrating these coordinate systems (Visium: hexagonal spot; MALDI-MSI: Cartesian) requires resampling or interpolation and resolution differences (Visium spots: diameter ∼55 µm with a center-to-center spacing of 100 µm; MALDI-MSI pixels: ∼60 µm) further complicate alignment. These disparities in geometry and resolution can lead to misalignment, loss of spatial detail, and challenges in downstream multimodal analyses. Our framework can identify similarities even with slight misalignment between modalities without the need for pixel-to-pixel co-registration.

A central methodological decision in this work was to avoid defining gene–lipid similarity using a single statistical metric. The primary reason for this is that spatial biological similarity is inherently multidimensional. Gene expression and lipid composition are measured using fundamentally different technologies that operate on different physical principles and exhibit distinct intensity distributions, spatial resolutions, and noise characteristics. As a result, no single metric is sufficient to characterize cross-modal spatial agreement robustly.

Therefore, we designed a MOS framework that integrates complementary spatial descriptors into a unified similarity score. Conceptually, this approach is analogous to how humans evaluate resemblance between related individuals [51], [52]. When we say that two siblings look alike, we are not basing that impression on a single trait – such as hair color or the shape of their eyes. Instead, the sense of resemblance emerges from the combined similarity of many features: facial structure, skin tone, proportions, and even subtle expressions (Figure 4). No single attribute is enough on its own. It is the consistent alignment across several independent traits that creates a convincing and reliable perception of similarity.

**Figure 4.**
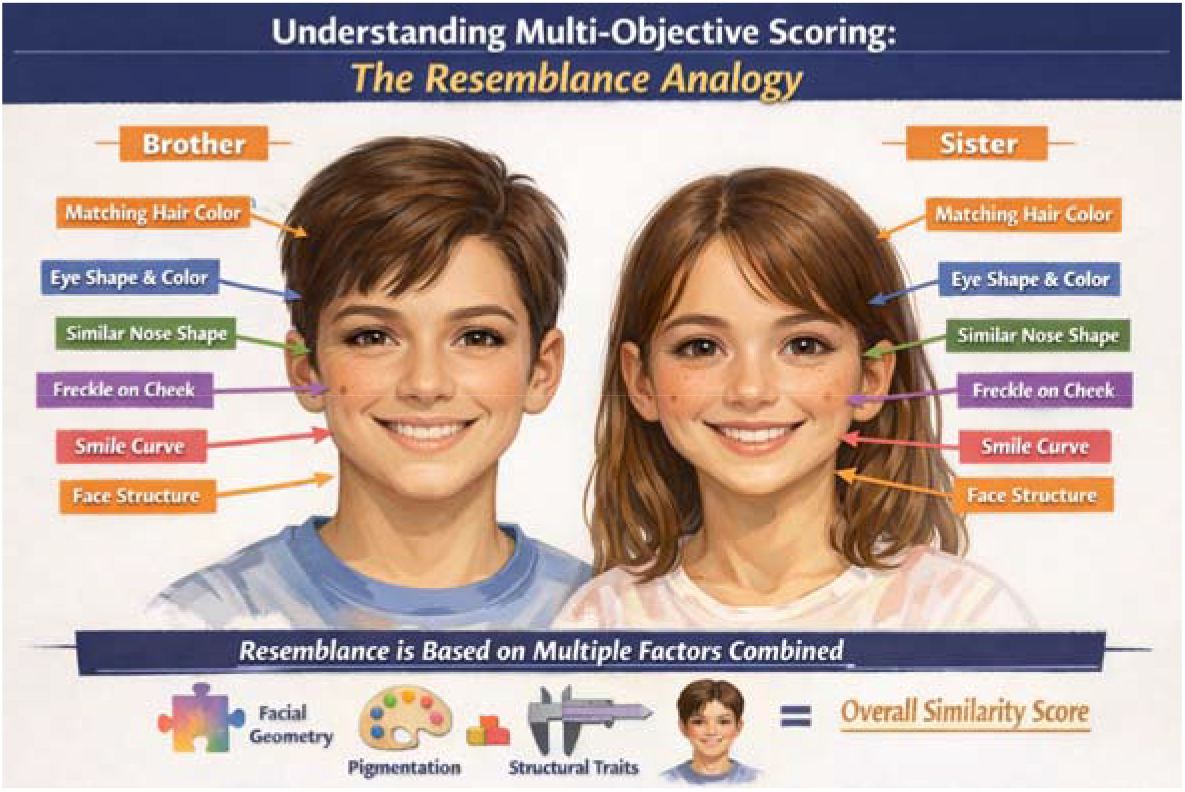
schematic demonstration of sibling resemblance^c^

The same principle applies to spatial molecular similarity. A gene and a lipid may share similar global intensity trends but differ in local clustering structure. Alternatively, they may exhibit similar clustering strength but be spatially shifted. If similarity were defined solely by a single metric – such as one Pearson correlation of intensity value – important structural differences could be overlooked, or false matches could be incorrectly accepted. For this reason, the similarity evaluation was decomposed into complementary components, each capturing a distinct dimension of spatial organization.

### Architecture and Mathematical Formula of Multi-Objective Scoring Framework

The MOS Framework was engineered as a robust framework for cross-modal spatial correlation. Unlike deep-learning approaches that require extensive training, this pipeline utilizes a heuristic-analytic architecture. The “match” between a target gene (G) and a candidate lipid (M) is quantified by a cross-modal similarity scoring engine (final combined score). This engine integrates direct spatial overlap with geometric descriptor similarity into a single scalar value (Figure 5).

**Figure 5.**
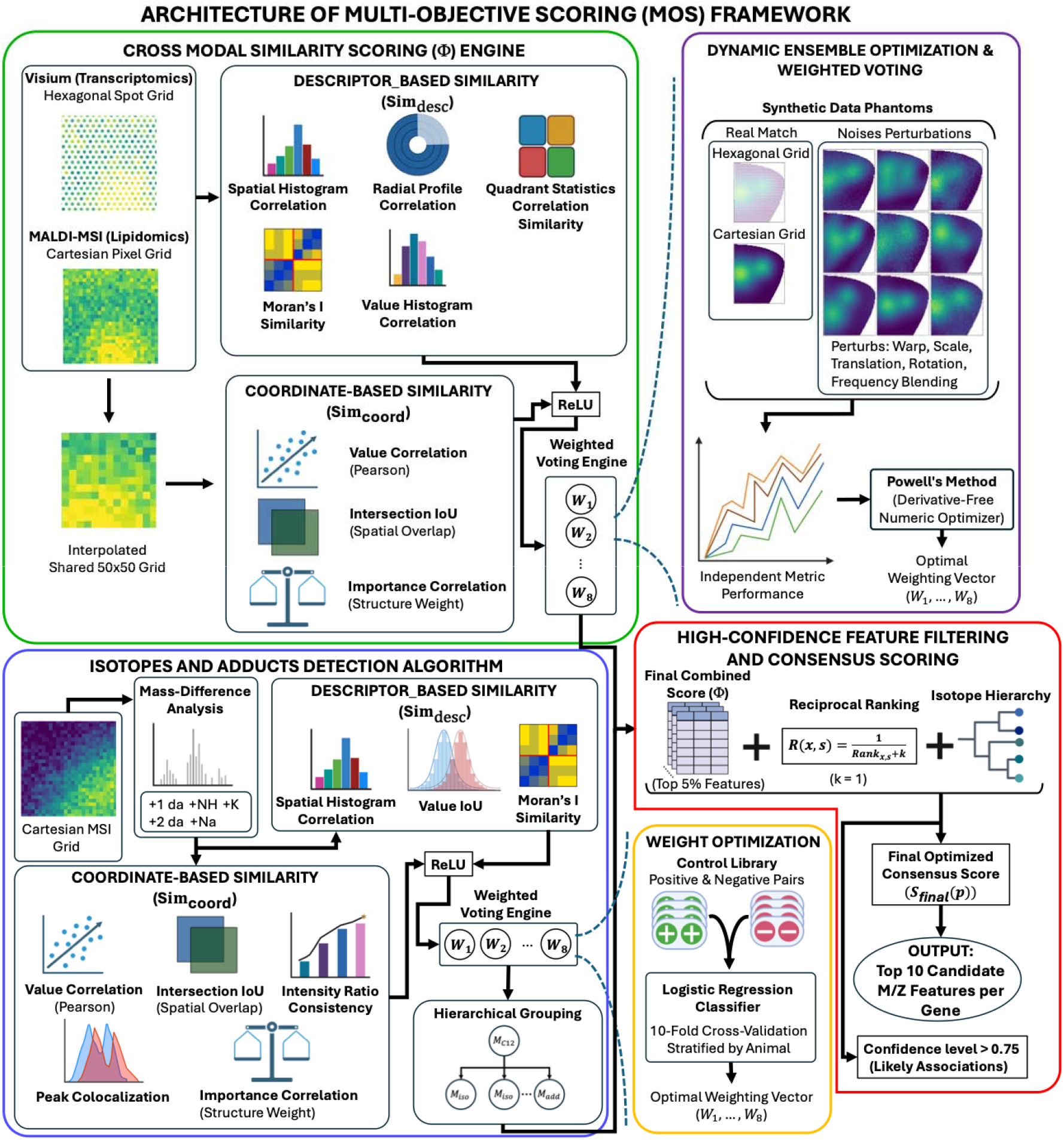
Schematic diagram of the MOS framework^d^

#### Component Metrics

Let *G*(*x,y*) represent the spatial pattern of a gene, and let *M*(*x,y*)represent the corresponding m/z. To quantify similarity, we divided the evaluation into two complementary components:

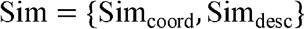

where coordinate-based similarity measures direct spatial alignment, and descriptor-based similarity evaluates higher-order structural relationships between the modalities.

#### Coordinate-Based Similarity (Sim _coord_)

All spatial maps were first geometrically ali gned a nd then interpolated onto a shared 50×50 grid to ensure a common spatial resolution. Let 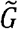 and 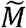 denote the interpolated gene and m/z grids, respectively.

##### a) Value Correlation

Spatial intensity similarity was quantified using Pearson’s correlation coefficient[53]:

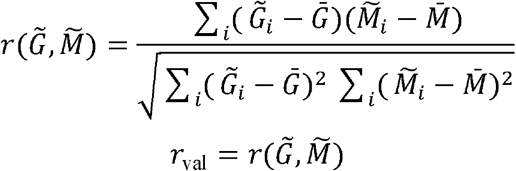

This metric measures the linear relationship of spatial intensity distributions, which means it measures how strongly the intensities of the two signals vary together across the tissue. Signals that increase and decrease in the same spatial regions produce higher similarity scores, while negative correlations are treated as non-similar relationships.

##### b) Importance Intersection over Union (IoU)

The importance map was computed for each signal by combining normalized intensity with local spatial variance among neighboring pixels. Two metrics were derived from these maps: Importance Intersection-over-Union and Importance Correlation. Let *I*_G_ and *I*_M_ represent normalized biological importance maps. Spatial overlap is defined as:

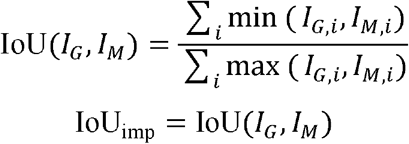

This metric quantifies the degree of spatial overlap of high-importance regions.

##### c) Importance Correlation

Pearson’s correlation coefficient is used to measure spatial similarity between the importance maps:

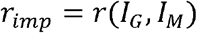

which evaluates the linear relationship in biologically weighted spatial structure.

Together, these metrics quantify pixel-level spatial correspondence after mild alignment and interpolation.

#### Descriptor-Based Similarity (Sim _desc_)

To capture structural similarity independent of exact pixel-to-pixel alignment, each spatial map is converted into a set of global descriptors. These descriptors summarize its distributional characteristics, geometric properties, and spatial autocorrelation patterns.

##### a) Spatial Histogram Correlation

Let *H*_*G*_ and *H*_*M*_ represent normalized 2D spatial histograms. Similarity is defined as:

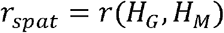

This metric compares coarse spatial distributions by dividing the tissue into grid regions and evaluating whether both signals show similar large-scale intensity patterns across those regions.

##### b) Radial Profile Correlation

Let *R*_*G*_ and *R*_*M*_ represent radial intensity profiles computed from the centroid outward. Similarity is defined as:

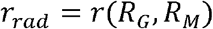

This metric measures how intensity changes from the center toward the periphery, enabling comparison of overall gradient patterns between the two maps.

##### c) Quadrant Statistics Correlation

The spatial domain is divided into directional subregions. Let *Q*_*G*_ and *Q*_*M*_ represent the concatenated quadrant-wise summary statistics from each map. Similarity is then computed as:

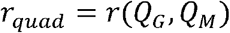

This metric assesses the degree of directional symmetry and regional heterogeneity shared between the two spatial patterns.

##### d) Moran’s I Similarity

Spatial autocorrelation is measured using Moran’s I [54], [55]:

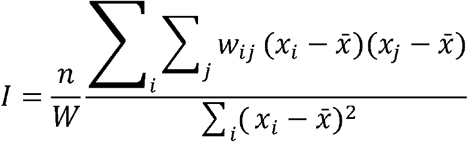

Where *w*_*ij*_ represents the spatial weights between locations *i* and *j*, and *W*=∑ _*i,j*_ *w*_*ij*_ is the total sum of weights.

In this implementation, the weight matrix *W* is constructed using a k-nearest neighbor (kNN) graph based on normalized spatial coordinates. The value of *k* was set to 6 for gene expression data and 8 for m/z data. This approach helps preserve local spatial structure while maintaining robustness to variations in sampling density.

Similarity between gene and m/z spatial autocorrelation is defined as:

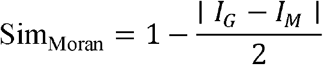

This formulation guarantees that the similarity score remains bounded within the interval [0,1]. Moran’s I spatial autocorrelation measures the tendency of neighboring pixels to exhibit similar intensities. The similarity between two signals is then defined by how closely their spatial autocorrelation values match.

##### e) Value Histogram Correlation

Let *V*_*G*_ and *V*_*M*_ represent normalized intensity histograms. Structural similarity in signal distribution is quantified via:

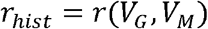

This metric captures how closely the two distributions are similar in terms of their overall shape and dynamic range, independent of spatial location.

#### Final Scoring Equation

The final combined score (Φ) is calculated as the sum of the weighted coordinate and descriptor scores. To ensure the model favors positive associations, correlation values are rectified using a max(*x*, 0) function where appropriate.

The total score Φ is defined as:

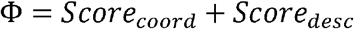

Breaking this down into the specific weights (*w*_1_ through *w*_8_) used in the pipeline:

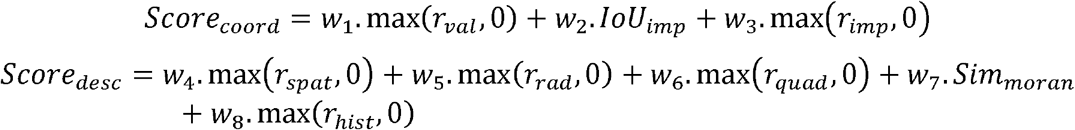

The weights were assigned to prioritize direct spatial overlap (*Score*_*coord*_) as the primary signal, while using the descriptor ensemble (*Score*_*desc*_) to validate the structural integrity of the match. The weights were calculated based on an optimization algorithm using synthetic data.

### Synthetic data

To evaluate the performance of the final combined score (Φ) framework and optimize its weight assignments (*w*_1_ through *w*_8_), a generative framework was developed to produce synthetic “digital phantoms.” These datasets are designed to replicate the biological variability and technical challenges of the spatial transcriptomics (RNA) and MSI. The simulation begins by defining a reproducible tissue morphology modeled after a murine brain hemisphere, constructed using a composite masking function that combines trigonometric and elliptical curves. To capture the physical acquisition differences between modalities, the tissue region was sampled using two distinct coordinate systems: a 100 µm hexagonal lattice for transcriptomic spots and a 60 µm Cartesian grid for mass spectrometry pixels. This variation in sampling density and geometry ensures that the synthetic data realistically captures the non-aligned nature of real-world spatial omics experiments.

Within this defined tissue region, the framework constructed 50 distinct spatial archetypes, ranging from simple linear gradients and periodic wave patterns to more biologically inspired structures such as cortical layers, which serve as the ground-truth (GT) signals. Each GT gene pattern is matched with a mathematically identical GT lipid pattern, establishing a known 1.0 correlation for validation purposes. To rigorously test the robustness of the scoring engine, a non-rigid distortion engine generates 10 “decoy” noise variants for every pattern. These decoys are produced through five independent spatial perturbations: sinusoidal coordinate warp (shape deformation), anisotropic scaling, center translation, rotational shifts, and frequency blending. The severity of each perturbation is set to 8 on a user-defined 1–20 scale. This controlled setup enables a systematic evaluation of the algorithm’s ability to distinguish true cross-modal biological associations from spatially coincident noise, providing the positive and negative control pairs required for the weight optimization process.

### Independent Metric Performance and Rotational Accuracy

The framework evaluated the robustness of the final combined score (Φ) engine by measuring the performance of each spatial metric separately across all ground-truth (GT) patterns and their associated orientations. In this process, gene expression data was systematically rotated, and the corresponding m/z matches were identified. The start, stop, and step angles are user-defined based on the orientation of the real data; in this study, they were set to 354°, 6° (±6°), and 2°, respectively. For every combination of gene expression, sample, and rotation angle, each of the eight scoring metrics – such as Value Correlation, Moran’s I Similarity, and Radial Profile Correlation – is treated as an independent classifier. By isolating these metrics, the analysis can determine which specific spatial descriptors are most resilient to the five types of spatial perturbations (warp, scale, translation, rotation, and frequency blending) introduced by the distortion engine.

At each orientation, the algorithm selects the m/z feature with the highest score for a given metric. A match is considered correct only if the top-scoring m/z corresponds to the pre-defined ground-truth lipid paired with the gene expression pattern. This approach makes it possible to generate detailed accuracy profiles for each metric, showing, for example, whether the Radial Profile metric better captures concentric ring structures, while Moran’s I performs more effectively for clustered “blob” patterns. The framework recorded these accuracies for every gene expression (phantom), sample, rotation, and scoring metric, and these results were subsequently used to inform and refine the weight assignments.

To maximize the predictive performance of the final combined score (Φ), a dynamic ensemble optimization strategy was implemented to determine the optimal weighting vector (w□, …, w□). Instead of relying on fixed weights, the framework employs a derivative-free numerical optimizer (Powell’s method [56], [57]) to maximize a global accuracy objective function within defined rotational windows. This approach allowed the model to adaptively prioritize different spatial descriptors based on their empirical performance within specific angular tolerances.

### Isotopes and Adducts Detection Algorithm

In many MALDI-MSI studies, isotopic peaks and adduct forms are removed during preprocessing to reduce redundancy in downstream analyses. In this framework, however, these related signals were intentionally retained and used as additional evidence to strengthen the MOS system. Because isotopes and adducts of the same molecule are expected to share highly similar spatial distributions, they provide an internal reference for evaluating the robustness of spatial similarity metrics.

To identify isotopes and adducts, a dedicated algorithm was developed that integrates two complementary components: (i) mass-difference analysis and (ii) spatial pattern similarity assessment. In the mass-difference step, candidate pairs are screened based on established isotopic spacing rules and commonly reported adduct mass shifts described in the literature. In the spatial component, candidate signals are evaluated to determine whether they exhibit consistent spatial organization across the tissue section. For spatial comparison, the same conceptual framework used in the cross-modal similarity scoring engine was adopted, with the simplification that all computations were performed directly within a Cartesian grid. In the gene-lipid matching pipeline, spatial alignment is more complex because the two modalities originate from different coordinate systems, requiring hexagonal-to-Cartesian interpolation and alignment that accounts for probable rotational differences. In contrast, isotope and adduct analysis was performed entirely within a single 60 µm Cartesian MSI grid. Because all signals originated from the same tissue section and coordinate system, no geometric transformation or rotational normalization was required. This allowed spatial similarity metrics to be computed directly across corresponding pixels without cross-modality alignment.

To evaluate spatial similarity between two m/z signals, the analysis was divided into two complementary components: coordinate-based similarity and descriptor-based similarity.

Coordinate-based similarity incorporates some metrics from the final combined score, including Value Correlation, Importance IoU, and Importance Correlation. It also uses two new metrics, including Intensity ratio consistency and Peak colocalization. Intensity ratio consistency assesses whether the relative intensity between the two signals stays consistent across spatial locations, as expected for isotopic peaks. Peak colocalization measures the spatial overlap of the highest-intensity pixels of each signal, indicating whether the strongest signal regions coincide within the tissue. The descriptor-based similarity incorporates Moran’s I, Spatial Histogram Correlation – from the final combined score, and value intersection-over-union. The value intersection-over-union metric evaluates the overall similarity of the raw intensity distributions across spatial locations, capturing similarity in signal magnitude independent of spatial structure. Together, these eight complementary metrics captured multiple aspects of spatial similarity expected for isotopes.

The weighting vector (w□, …, w□) for the spatial metrics was refined using a supervised machine learning model (Logistic Regression) based on previously validated isotope and adduct relationships. For each sample, a control library consisting of 20 positive control pairs and 18 negative control pairs (lipid signals with clearly distinct spatial distributions) was constructed. The positive pairs were detected based on mass difference, intensity ratio, and overall pattern [58], [59]. The complete list of positive and negative controls is provided in Table 3 and Table 4. The same control library was applied consistently across all samples to ensure comparability.

**Table 3.**
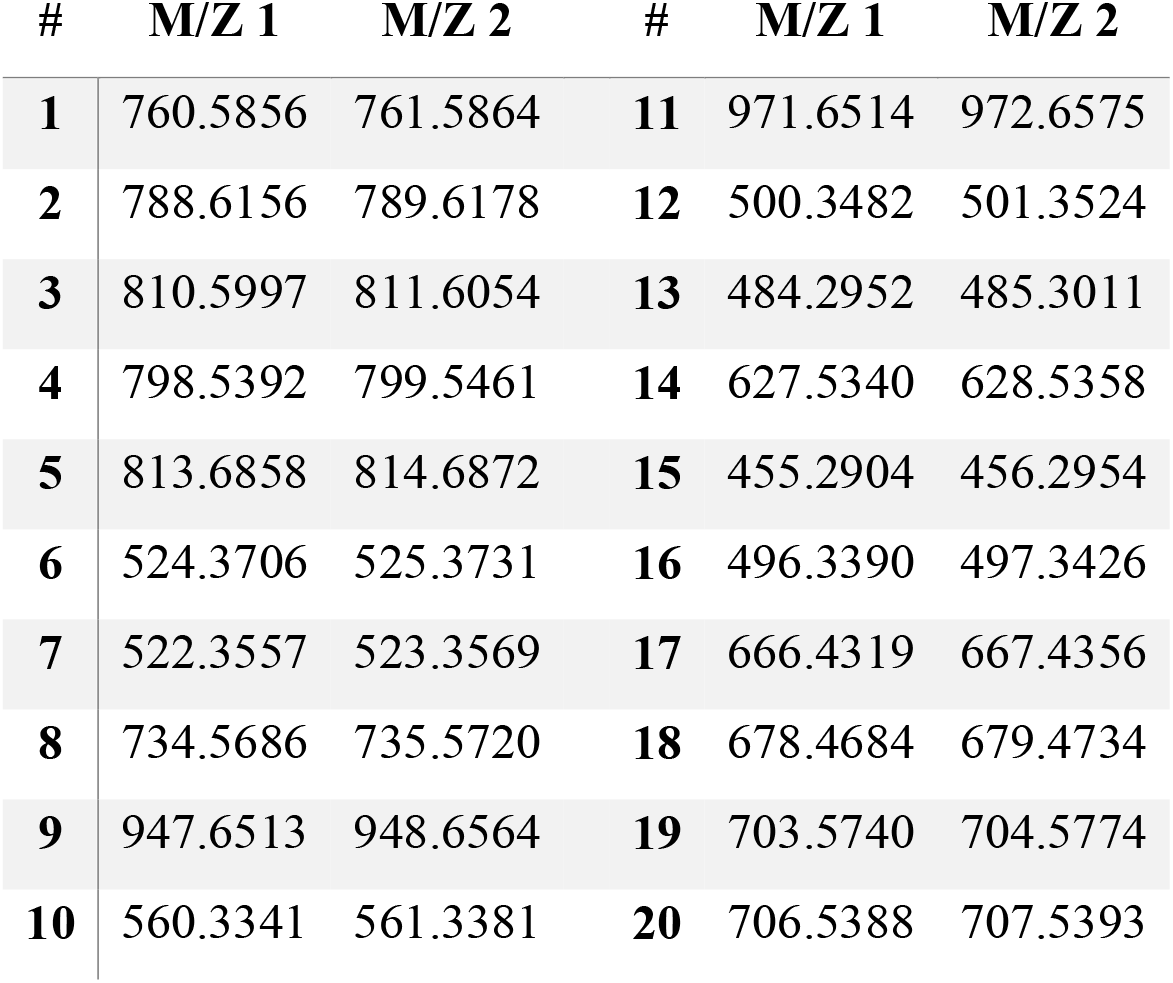
Positive Control Pairs.

| # | M/Z 1 | M/Z 2 | # | M/Z 1 | M/Z 2 |
| --- | --- | --- | --- | --- | --- |
| 1 | 760.5856 | 761.5864 | 11 | 971.6514 | 972.6575 |
| 2 | 788.6156 | 789.6178 | 12 | 500.3482 | 501.3524 |
| 3 | 810.5997 | 811.6054 | 13 | 484.2952 | 485.3011 |
| 4 | 798.5392 | 799.5461 | 14 | 627.5340 | 628.5358 |
| 5 | 813.6858 | 814.6872 | 15 | 455.2904 | 456.2954 |
| 6 | 524.3706 | 525.3731 | 16 | 496.3390 | 497.3426 |
| 7 | 522.3557 | 523.3569 | 17 | 666.4319 | 667.4356 |
| 8 | 734.5686 | 735.5720 | 18 | 678.4684 | 679.4734 |
| 9 | 947.6513 | 948.6564 | 19 | 703.5740 | 704.5774 |
| 10 | 560.3341 | 561.3381 | 20 | 706.5388 | 707.5393 |

**Table 4.**
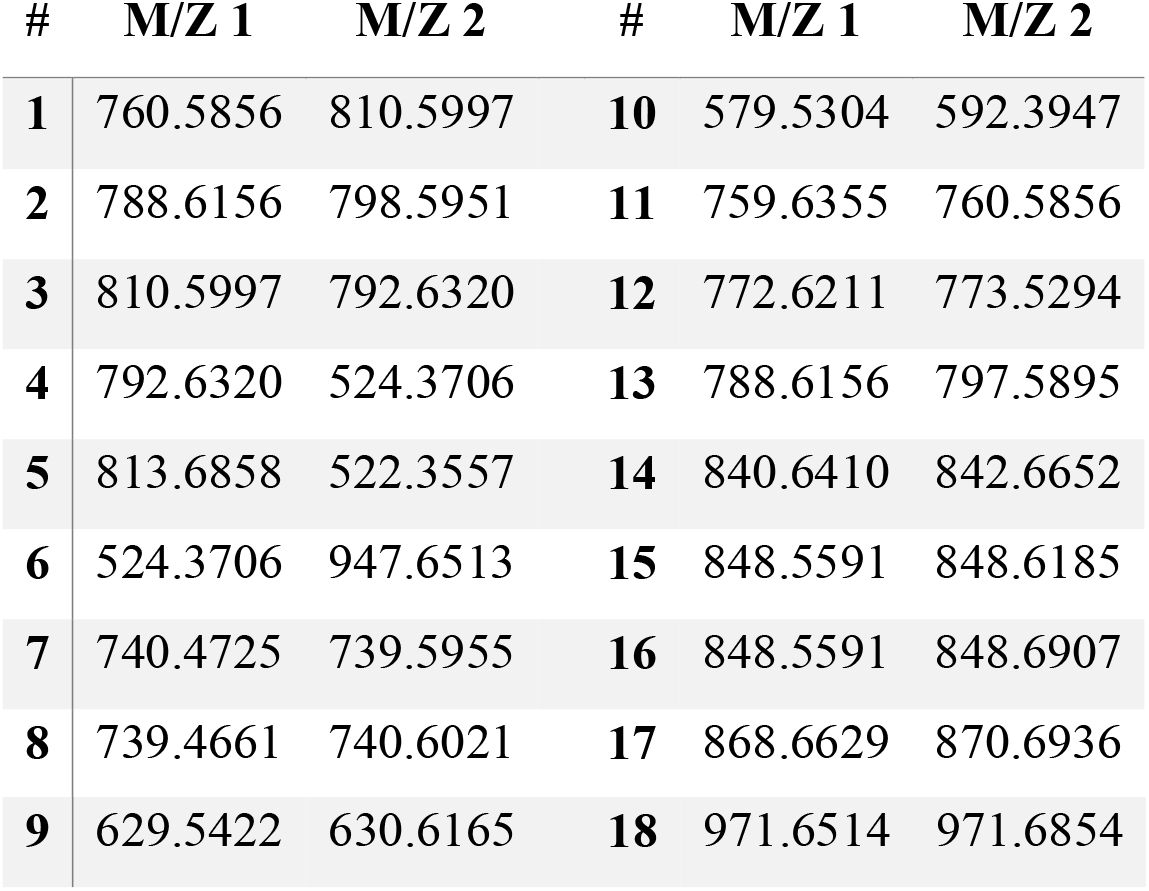
Negative Control Pairs (Spatially Distinct Lipid Signals)

In the Logistic Regression classifier, model training employed 10-fold cross-validation, stratified by animal, to ensure that learned weights generalized across biological variability among the young control (YC), young Alzheimer’s disease (YAD), aged control (AC), and aged Alzheimer’s disease (AAD) groups. This design prevented overfitting to tissue-specific spatial characteristics and ensured robustness to inter-animal heterogeneity.

The purpose of this procedure was to identify metric weights that maximize discrimination between true isotope/adduct pairs and spatially unrelated signals that may appear close in mass. Model performance was evaluated using the standard error (SE) of metric coefficients across folds. Metrics demonstrating consistently low SE values were prioritized due to their stability across tissue samples and biological conditions.

#### Isotopic and Adduct Hierarchy Mapping

After determining the weights, the algorithm was employed to identify and merge chemically related m/z features based on their spatial co-localization. Let’s M_C12_ denote the most abundant isotope of an ion (e.g. ^12^C) and M_iso/add_ its other isotopes or adducts (e.g. ^13^C).

- Pattern Matching: The algorithm searched for five predefined mass shifts accounting for isotopes and adducts: +1 da, +2 da, +NH, +Na, and +K.
- Frequency Filtering: A candidate relationship was only validated if it surpassed a 60% score threshold in at least 12 out of the 16 samples (>=75% of the cohort).
- Hierarchical Grouping: For each M_C12_ m/z feature, the algorithm identified the closest match within a 0.01 Da tolerance. Validated features were organized into a strict M_C12_-M_iso/add_ hierarchy, limiting each M_C12_ to a maximum of five M_iso/add_ to maintain chemical plausibility.

### High-Confidence Feature Filtering

The MOS framework was developed to systematically evaluate and rank each m/z feature within the dataset, assigning a performance score to all 528 unique values for every gene under investigation. A key component of the framework is the estimation of a confidence level, which is defined by the frequency with which an m/z feature is detected across the murine cohort. This strategy was designed to balance sensitivity and specificity. Wider inclusion thresholds may increase the risk of false positives and artificially raised confidence estimates, whereas overly restrictive thresholds may exclude true positives and potentially filter out biologically meaningful lipid associations.

Because the optimal threshold depends on factors, such as the number of m/z features, spatial patterns, sample quality, and overall data quality, only the top 5% of features were retained in this study. From the initial pool of 528 unique m/z features ranked in each murine brain sample using the final combined score (Φ), the highest 26 features were selected as primary candidates for the gene-to-lipid matching process.

### Consensus Scoring and Chemical Synergy

For each gene, the retained top 5% of m/z features ranked using the final combined score (Φ) were further evaluated to determine a consensus ranking across all samples. To integrate evidence across samples while accounting for chemical structure, a hierarchical rank-based scoring framework was developed that combines each M_C12_ m/z feature with its validated isotopes or adducts.

#### Rank-Based Contribution

Within each sample, detected m/z features were ranked based on their Φ. To transform rank into a quantitative contribution, we applied a modified Reciprocal Rank formulation:

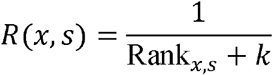

where:

Rank_*x,s*_ represents the rank of m/z feature *x* within sample *s*.

The parameter *k* is a smoothing constant that controls how strongly rank influences the score *R(x, s)*. In this study, it was set to 1 to give greater emphasis to higher-ranked features. This formulation ensures that top-ranked features contribute more significantly to the final score, while still allowing a gradual decrease in influence for lower-ranked features.

#### Hierarchical Integration of Isotopes and Adducts

Each M_C12_ m/z feature may have validated isotopes or adducts, represented as 𝒜(*p*), defined from the determined M_C12_-M_iso/add_ hierarchy file.

For each sample *s*, the contribution of the M_C12_ feature, represented as *S*_*p,s*_, is calculated as follows:

- *Case 1: M*_*C12*_ *m/z detected* If the M_C12_ m/z feature *p* is detected in the sample *s*, its contribution is:

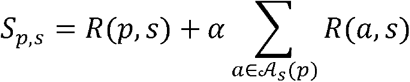

where: 𝒜_*s*_ (*p*) ⊆ 𝒜(*p*) are isotopes or adducts of *p* detected in the sample *s*, The parameter α represents the synergy weight. In this study, α was set to 0.2 to reduce the influence of isotopes and adducts. Lower α ensures that the isotopes and adducts combined contribution does not dominate the overall ranking or mask other high-ranking m/z features within the top 10. Based on this formulation, the M_C12_ feature contributes fully based on its rank, and co-detected isotopes or adducts provide additional, proportionally weighted reinforcement.
- *Case 2: M*_*C12*_ *absent, isotopes/adducts detected* If the M_C12_ m/z feature is not detected in sample *s*, but one or more of its isotopes or adducts are present, these related signals are treated as supporting chemical evidence. In this case, the per-sample contribution is calculated as:

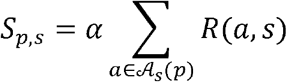

Accordingly, indirect chemical support contributes to the M_C12_ feature’s score, but at a reduced weight to avoid overestimating its importance.
- *Case 3: No detection* If neither the M_C12_ nor any of its isotopes or adducts is detected:

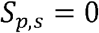

#### Final Optimized Consensus Score

The final optimized consensus score for a M_C12_ m/z feature *p* across all samples 𝒮 is defined as:

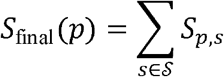

Expanded form:

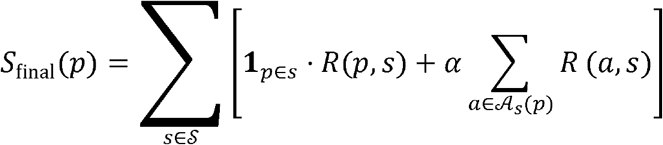

Where **1**_*p*∈*s*_ equals 1 if the M_C12_ m/z is detected in the sample *s*, and 0 otherwise.

#### Confidence Score

The confidence level is defined as the proportion of samples in which either the M_C12_ or at least one of its isotopes/adducts is detected:

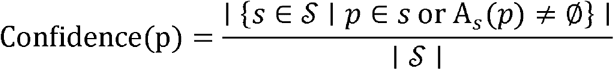

The MOS framework reports the top 10 candidate m/z features for each gene based on their final score, *S*_final_ (*p*), and assigns a confidence level to each candidate. Candidates with a confidence level exceeding 0.75 are considered more likely to be associated with the expressed gene.

## Results and Discussion

After the preprocessing stage of the MALDI-MSI data, 528 peaks were retained for each of the 16 samples for downstream analysis. Subsequently, two independent optimization algorithms were applied to determine metric weights for two distinct analytical objectives. The first optimization procedure calibrated the weights for the final combined spatial similarity score (Φ) used in gene-to-MSI pattern matching. The second optimization procedure determined the weights for the isotope and adduct detection algorithm, which integrates spatial similarity metrics within a classification framework. Although both approaches rely on spatial similarity analysis, they employ partially different metrics and descriptor implementations, and they were optimized using separate algorithms tailored to their respective objectives. This independent design ensures fair evaluation and robust performance for each task.

Table 5 reports the optimized relative contribution of the eight spatial similarity metrics used in the final combined score (Φ), which integrates coordinate-based overlap and structural descriptor similarity within the rotation-based pattern matching framework. The weights were calculated using synthetic data consisting of 50 structured spatial patterns and 500 noise patterns per each of the 16 samples, ensuring robust calibration under controlled ground-truth conditions. The Powell’s method optimization algorithm assigned the highest weights to Value_Hist_Corr (0.1850) and Radial_Corr (0.1814), indicating that similarity in intensity distribution and radial spatial organization provides the strongest evidence for true pattern correspondence. Importance_Correlation (0.1356) and Quadrant_Corr (0.1133) also contribute substantially, highlighting the importance of localized structural agreement across spatial regions. In contrast, Value_Correlation (0.0832) and Morans_Sim (0.0914) received lower weights, suggesting that global intensity similarity or spatial autocorrelation alone is less informative than the integrated set of structured spatial descriptors. Importantly, the optimized weighted combination achieved an average accuracy of 96.14% in the 354°–6° region, demonstrating that integrating complementary spatial features significantly enhances robustness and discriminative performance. These results confirm that the multi-component formulation of Φ – combining spatial overlap with descriptor validation – provides a reliable and empirically supported strategy for identifying biologically meaningful peak relationships in MALDI-MSI data.

**Table 5.** Optimized normalized weights of the eight spatial similarity metrics contributing to the final combined scoring function (Φ), derived through data-driven calibration using synthetic training data.

| **Metric** | **Optimal Weight** |
| --- | --- |
| Value_Correlation | 0.0832 |
| Importance_IoU | 0.1102 |
| Importance_Correlation | 0.1356 |
| Value_Hist_Corr | 0.1850 |
| Spatial_Hist_Corr | 0.1000 |
| Radial_Corr | 0.1814 |
| Quadrant_Corr | 0.1133 |
| Morans_Sim | 0.0914 |

Table 6 summarizes the calibrated contribution of eight spatial similarity metrics used to determine whether two MALDI-MSI peaks exhibit similar spatial patterns. These weights were estimated using a 10-fold cross-validated logistic regression model trained on positive and negative control *m/z* pairs extracted from 16 MSI brain samples.

**Table 6.** Mean normalized weights (± SD and SE) of spatial similarity metrics used in the pattern-similarity model for identifying isotopes and adducts among MALDI-MSI peaks, evaluated using 10-fold cross-validation.

| Metric Name | Weight (0–1) | SD | Std. Error (SE) |
| --- | --- | --- | --- |
| Intensity Ratio Consistency | 0.2231 | 0.0032 | 0.0010 |
| Value Correlation | 0.0414 | 0.0049 | 0.0016 |
| Peak Colocalization | 0.0850 | 0.0069 | 0.0022 |
| Importance IoU | 0.1226 | 0.0038 | 0.0012 |
| Importance Correlation | 0.1948 | 0.0030 | 0.0009 |
| Spatial Histogram Correlation | 0.1111 | 0.0056 | 0.0018 |
| Value IoU | 0.0508 | 0.0057 | 0.0018 |
| Moran's Similarity | 0.1713 | 0.0023 | 0.0007 |
| **TOTAL (Sum)** | **1.0000** |  |  |

As described in the Methods section, to ensure robustness and minimize overfitting, model performance was evaluated using 10-fold cross-validation grouped by animal sample. In each fold, data from specific animals were used for training, while data from previously unseen animals were reserved for testing. This animal-level separation ensures that the model generalizes across biological replicates rather than memorizing sample-specific patterns. The resulting mean AUC of 0.9675 demonstrates excellent discrimination between positive and negative peak relationships, confirming that the combined spatial similarity metrics reliably capture biologically meaningful ion co-localization.

The Weight (0–1) column represents the average normalized contribution of each metric across all ten folds. Because the logistic regression coefficients were normalized to sum to one, these values reflect the relative importance of each feature in the final calibrated model. Higher weights indicate greater influence in distinguishing true peak associations.

Among the metrics, intensity ratio consistency (0.2231) shows the highest contribution, indicating that consistent intensity relationships between ions across spatial pixels are the strongest indicator of true peak associations. Spatial structure metrics also contribute substantially, particularly importance correlation (0.1948), Moran’s similarity (0.1713), and Importance IoU (0.1226), highlighting shared biologically important regions, spatial autocorrelation, and the importance of spatial overlap. Moderate contribution is observed for spatial histogram correlation (0.1111), while peak colocalization (0.0850) has a smaller influence. In contrast, value IoU (0.0508) and value correlation (0.0414) contribute minimally, suggesting that global intensity correlation alone is not a strong indicator of true MSI peak relationships compared with spatially informed descriptors.

After applying the weights in the isotope and adduct detection algorithm, containing mass-difference analysis and spatial pattern similarity assessment metrics, and using it on all data, 136 peaks were identified as having at least one additional isotope or adduct. Subsequently, MOS introduced 10 candidates for each expressed gene by incorporating 528 m/z peaks for each sample, isotopes/adducts hierarchy, and 8 selected expressed genes (Figure 6).

**Figure 6.**
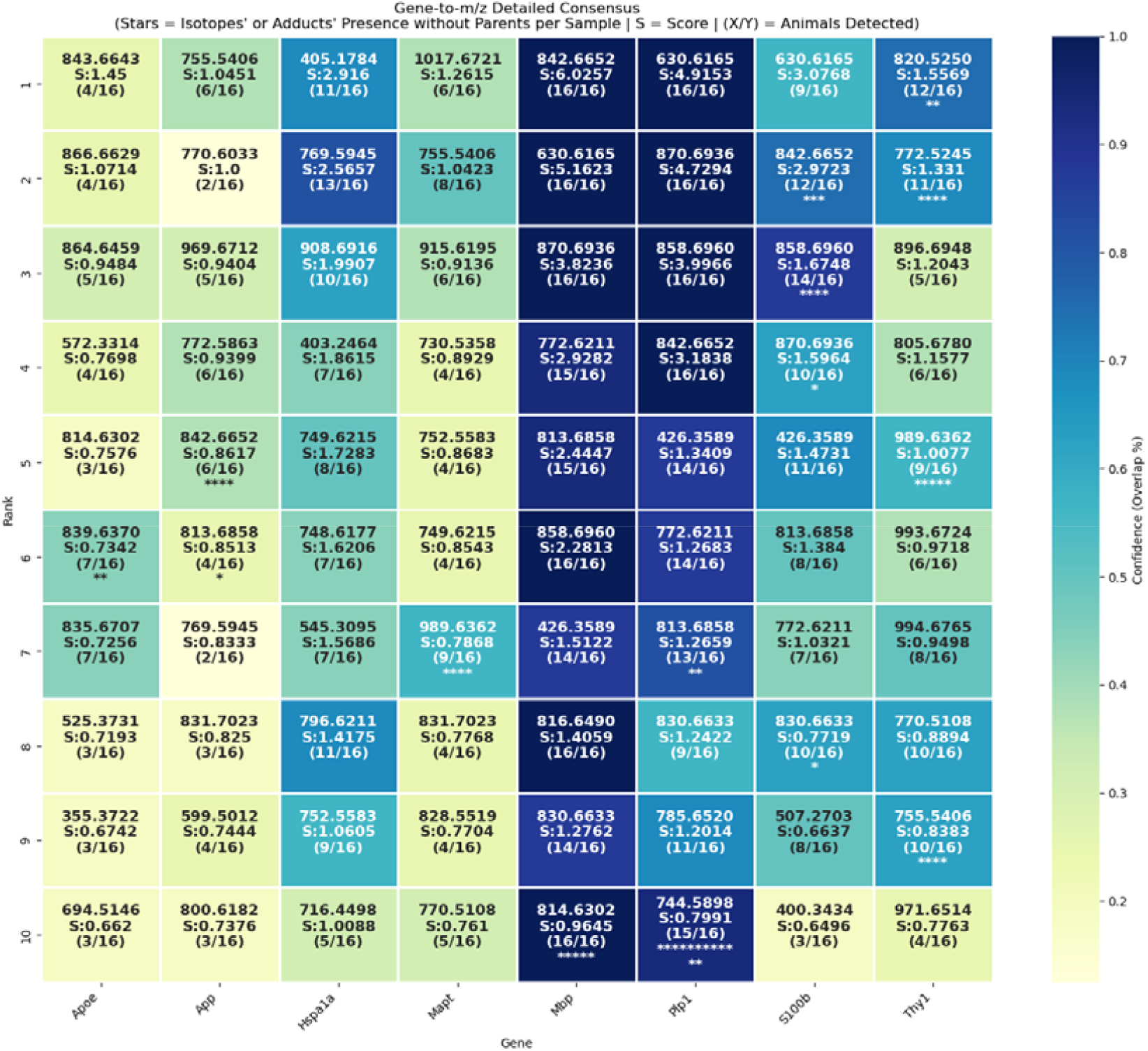
Gene–m/z consensus heatmap. Columns represent genes and rows show the top 10 ranked m/z candidates. Color indicates confidence (fraction of animals where the feature was detected in the top 5%). Each cell reports the m/z value, consensus score (S), and detection frequency (X/Y). Asterisks (*) denote samples where isotopic or adduct peaks were detected without the M_C12_ peak.

Based on the results (Figure 6) and considering the confidence score (>=0.75), the most robust associations were observed for the myelin-related genes *Mbp* and *Plp1*[60], [61]. Several m/z features linked to these genes were detected in nearly all animals, including 630.6165, 870.6936, and 842.6652, each appearing in all 16 samples. The high reproducibility and strong consensus scores of these peaks suggest that they represent stable analyte signatures associated with myelin-rich regions of the brain. Both *Mbp* and *Plp1* encode structural components of the myelin sheath produced by oligodendrocytes [60], [61], and their consistent association with the same analyte features indicates that these peaks may correspond to lipid species involved in myelin structure or maintenance.

Interestingly, one of the most recurrent features across multiple genes was m/z 842.6652, which appeared among the top-ranked peaks for *Mbp, Plp1*, and the astrocyte-associated gene *S100b* [62]. This peak was detected in all animals for *Mbp* and *Plp1* and in the majority of samples for *S100b*. The S100B protein is widely used as a molecular marker of astrocytes and glial activity in the central nervous system, although it can also be expressed in oligodendrocytes during development [62], [63], [64]. The repeated detection of the same m/z feature across these genes indicates a strong spatial similarity (overlap) between the gene-expression patterns and the analyte distributions across samples, suggesting overlapping molecular signatures in regions enriched with glial cells and myelinated structures.

Figure 7 shows the spatial heatmap of *Mbp* expression across all samples. The expression pattern follows the expected anatomical distribution of myelinated white matter tracts in the mouse brain. The corresponding MALDI-MSI distribution of analyte m/z 842.66 (Figure 8) displays a highly similar spatial pattern after TIC normalization and removal of background and noise pixels. Both maps show strong signal enrichment along white matter structures and reduced signal in cortical gray matter regions [65]. The close spatial similarity between the transcriptomic signal and the MSI analyte suggests that the MOS framework can successfully identify analyte features associated with myelin-related molecular patterns.

**Figure 7.**
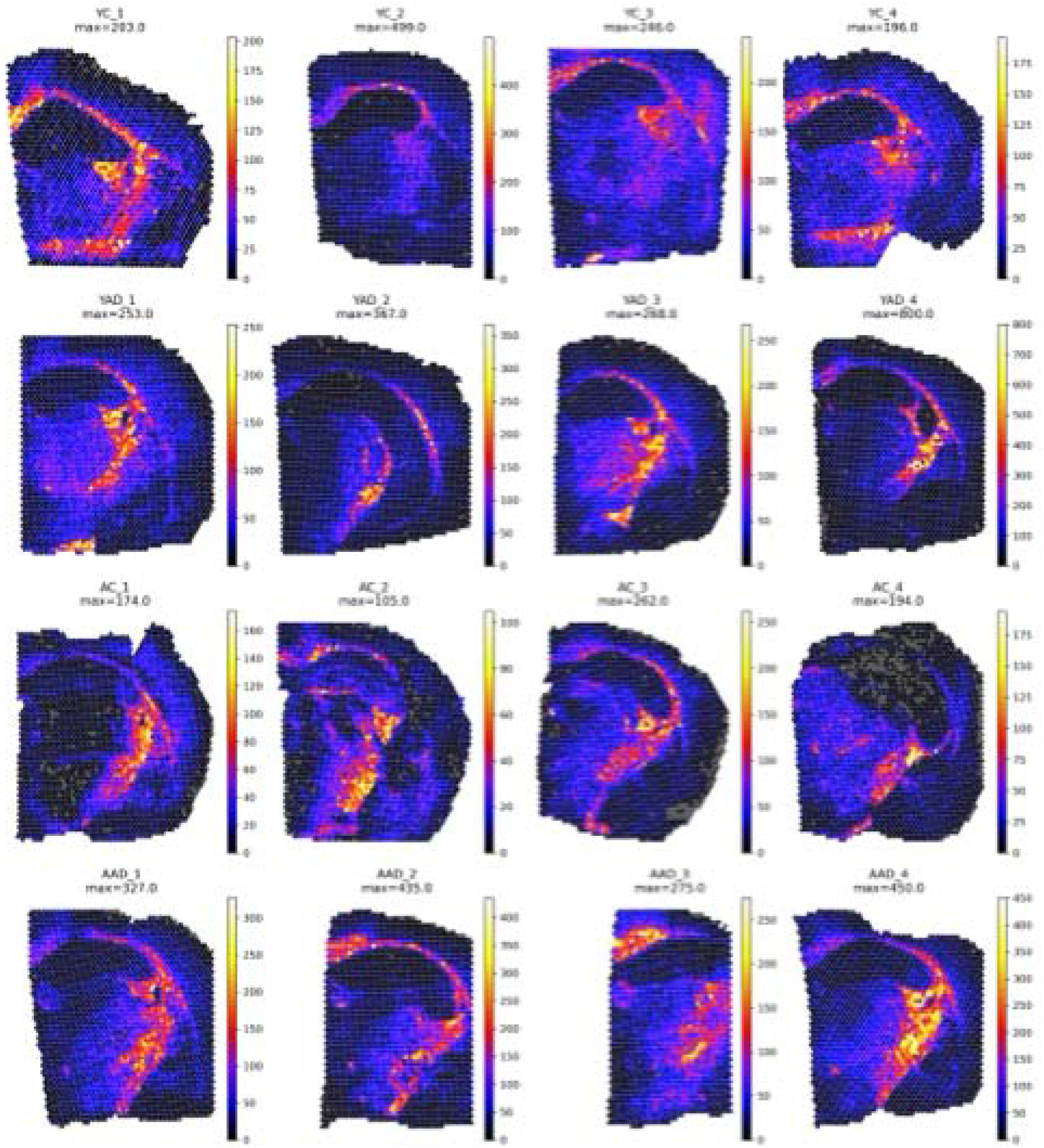
Spatial heatmap of *Mbp* expression across all samples; spots enlarged for easier visualization

**Figure 8.**
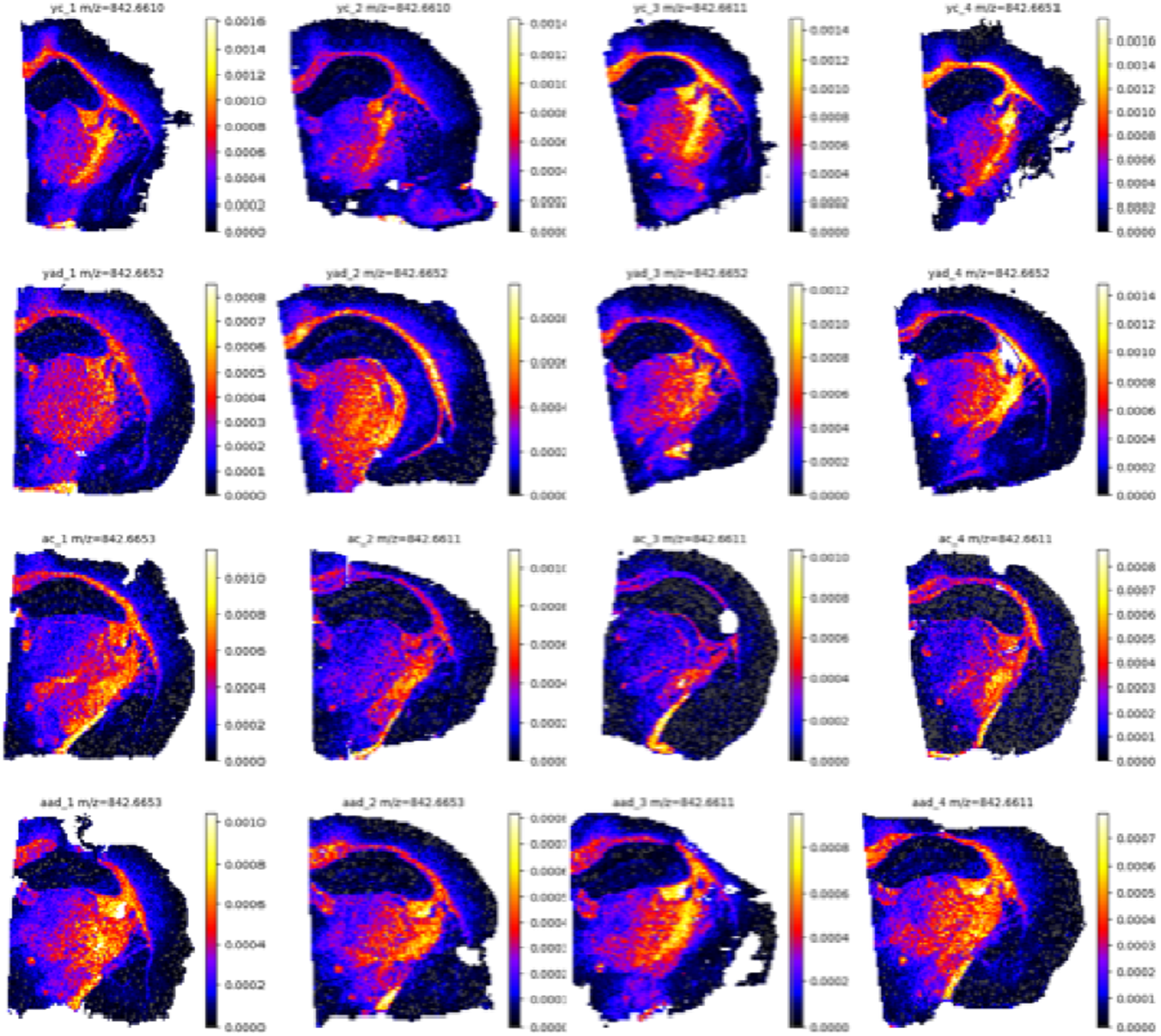
TIC-normalized spatial distribution heatmaps of analyte 842.6652 across all half-brain samples with filtering of background, probable contamination, or noise pixels

Figure 9 further demonstrates this relationship by comparing the spatial distribution of *Mbp* with the top ten candidate m/z features with high confidence level proposed by the MOS framework in a representative sample. All candidate features exhibit spatial distributions that closely resemble the *Mbp* expression map, indicating that the algorithm consistently identifies analytes localized to white matter regions. This result supports the robustness of the MOS scoring system when the gene expression pattern is spatially well-defined and densely sampled. The similarity among multiple candidate analytes may reflect the presence of several lipid species associated with myelin sheaths, which are known to be abundant in white matter and detectable by MALDI-MSI.

**Figure 9.**
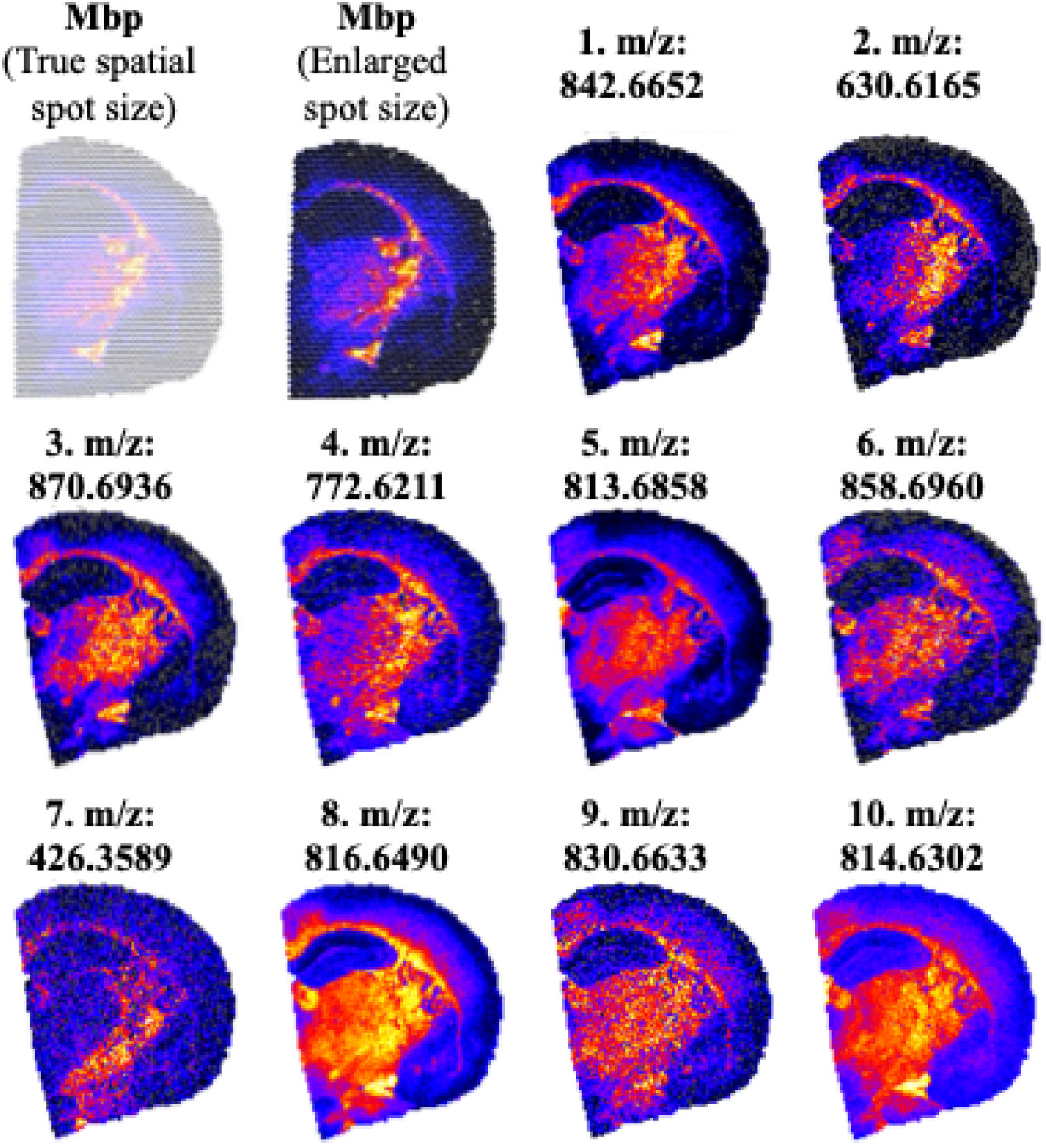
Spatial distribution heatmap of *Mbp* across a representative sample, shown with both true spatial spot size and enlarged spots for visualization, alongside the distributions of the 10 matched m/z candidate features identified using the MOS framework. Numbers indicate the ranking of the candidate features.

Figure 10 and Figure 11 illustrate a case in which only partial spatial similarity was observed between transcriptomic and MSI data. The spatial heatmap of *Thy1* expression (Figure 10) shows enrichment in cortical and hippocampal regions, particularly in AD samples, while the corresponding MSI analyte at m/z 820.52 (Figure 11) exhibits a broadly similar but imperfect spatial distribution. The discrepancy appears to be largely driven by artifacts at tissue boundaries, as the most prominent signal regions in both spatial maps are located near or directly along the tissue borders.

**Figure 10.**
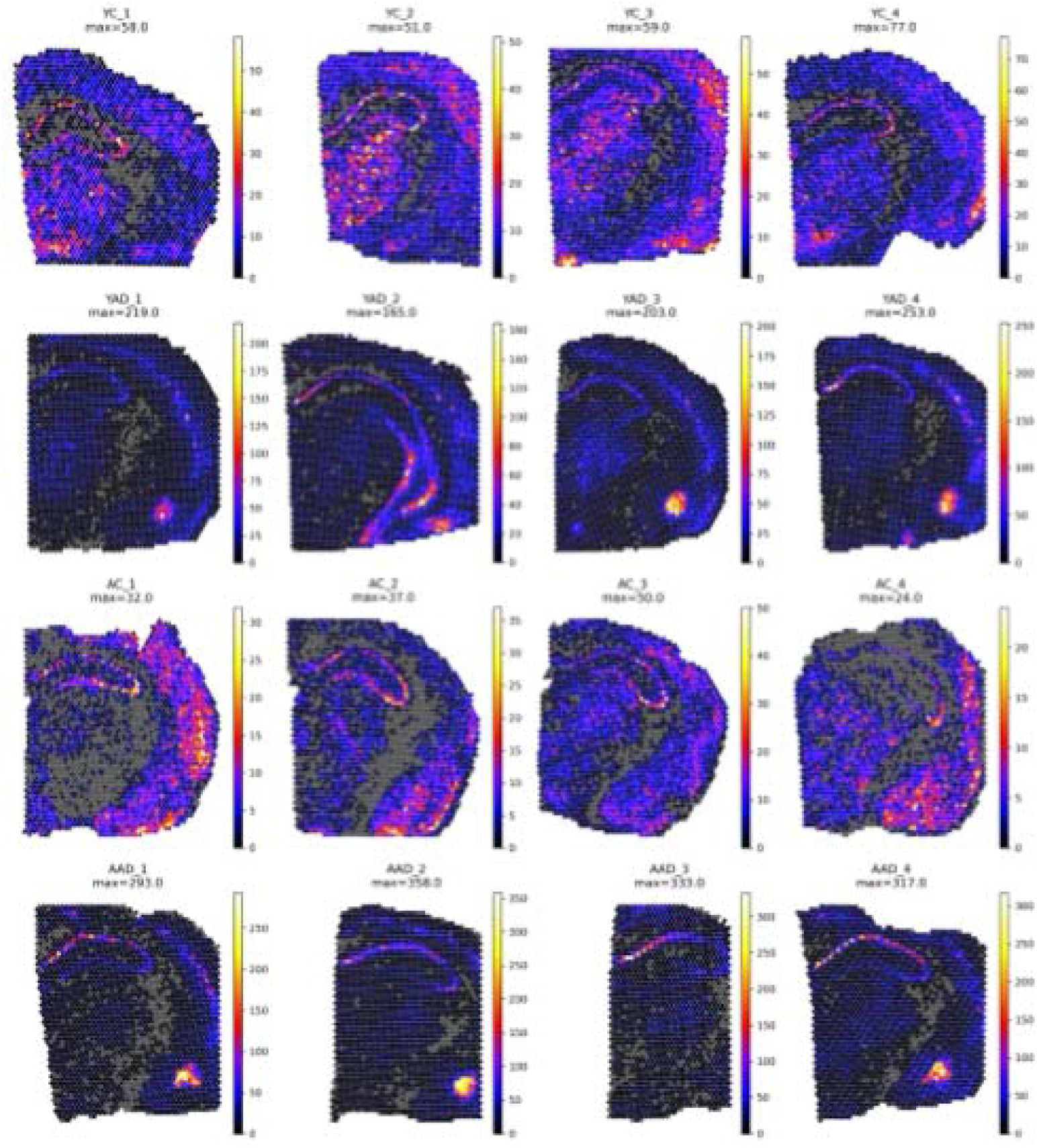
Spatial distribution heatmap of *Thy1* expression across all samples; spots enlarged for easier visualization

**Figure 11.**
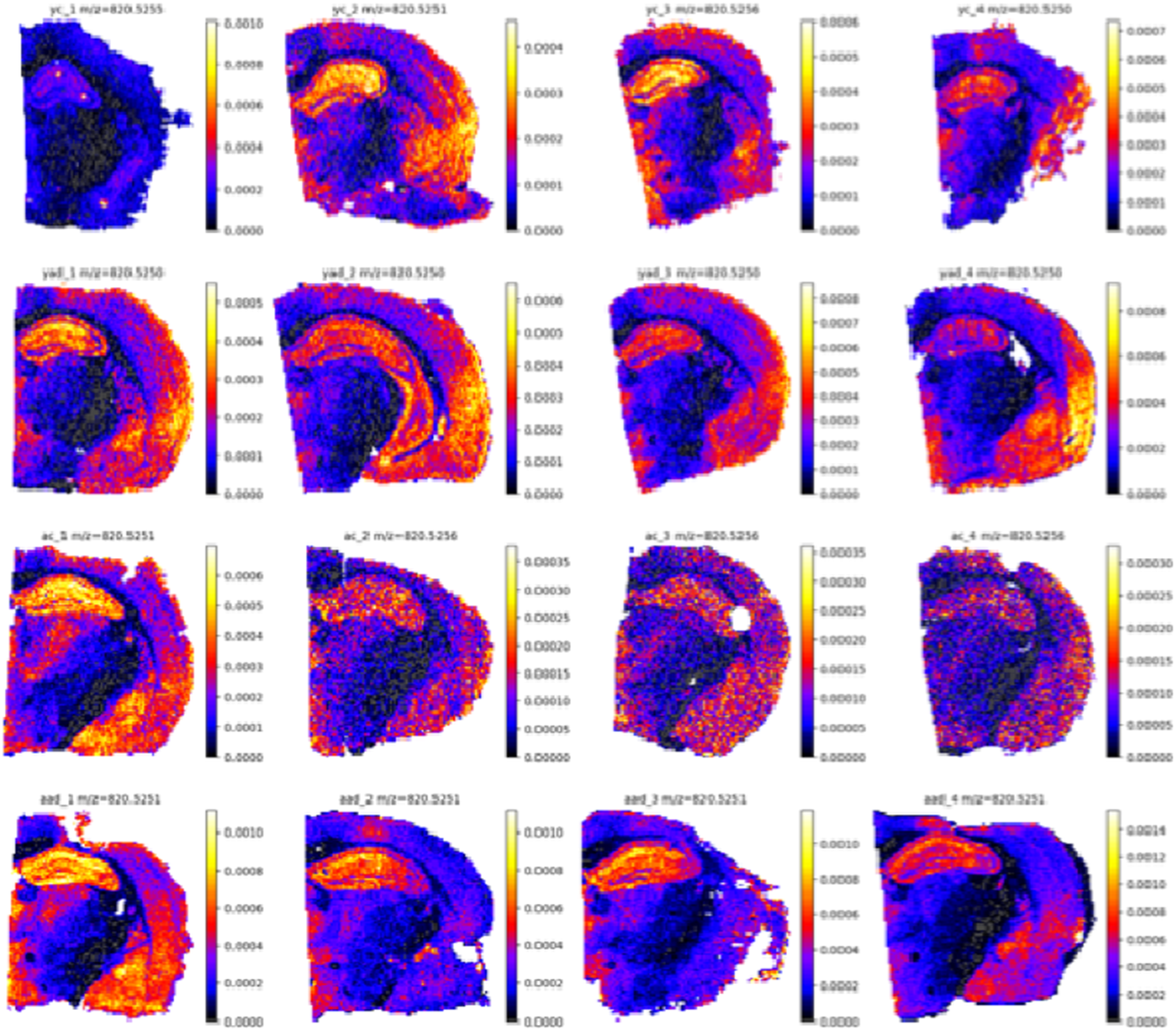
TIC-normalized spatial distribution heatmaps of analyte 820.52 across all half-brain samples with filtering of background, probable contamination, or noise pixels

In both modalities, edge regions are prone to distortions caused by issues during freezing, sectioning, and mounting. Additional inconsistencies in spatial transcriptomics data can be due to an incomplete capture or misaligned spots at the margins. As a result, some transcriptomic spots may fall partially outside the actual tissue region, while MSI pixels located near tissue edges may experience altered ionization efficiency or matrix-related effects. These factors can introduce spatial inconsistencies that reduce the accuracy of similarity assessment between the gene and analyte maps, as seen in the *Mbp* and *Thy1* cases. In the *Mbp* example, the most informative regions of expression were located closer to the center of the tissue section, where the data are less affected by edge-related distortions. In contrast, the prominent regions of *Thy1* expression and the corresponding MSI signal are positioned closer to the tissue boundaries, making them more susceptible to sample preparation artifacts and spatial misalignment. This difference likely contributes to the lower spatial agreement observed in the *Thy1* case compared with the *Mbp*-associated analytes.

Figure 12 presents an example highlighting a limitation of the MOS framework when gene expression data are sparse. The spatial heatmap of *Hspa1a* shows relatively few detected spots across the tissue section, resulting in a highly sparse expression map. Because the MOS framework assumes that available gene expression signals correspond to meaningful spatial information, the limited number of gene spots can artificially increase similarity scores across many MSI features. Consequently, the framework may assign a relatively high score to m/z value, such as the proposed candidate at m/z 769.59, even when a clear spatial correspondence is not evident. This issue suggests that gene quality-control and filtering steps as well as PCA and finding HVGs (highly variable genes) should be applied prior to the MOS analysis to remove sparsely expressed genes or low-confidence spots. By implementing such preprocessing steps in the pipeline, genes with insufficient spatial information will likely be excluded and will not appear among the proposed gene–lipid associations.

**Figure 12.**
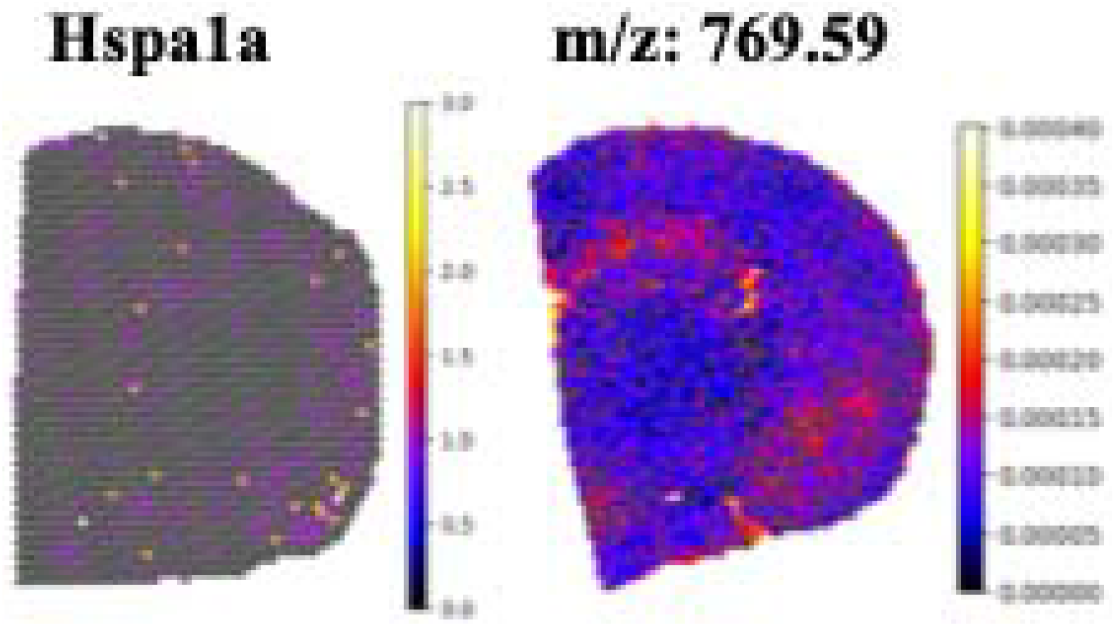
Spatial distribution heatmap of *Hspa1a* expression across a representative sample (left), with spots enlarged for clearer visualization, and the spatial distribution heatmap of its proposed candidate analyte feature m/z 769.59 (right), identified using the MOS framework.

## Conclusion

Overall, these results demonstrate that the MOS framework can effectively identify MALDI-MSI analyte features whose spatial distributions correspond to transcriptomic patterns. The optimized multi-metric scoring approach showed strong performance in both synthetic validation and experimental data, indicating that combining complementary spatial descriptors improves the reliability of pattern matching. In the biological examples, consistent associations were observed for the myelin-related genes *Mbp* and *Plp1*, whose expression patterns closely matched several analyte features, particularly m/z 842.6652, suggesting that these peaks may reflect lipid species associated with myelin-rich brain regions.

More broadly, the MOS framework is designed as a hypothesis-generation tool that examines large-scale spatial relationships while preserving structural details. The concept of a consensus score can be extended to other pattern-matching and similarity analyses, and its metric components could be adapted or replaced with alternative statistical, machine-learning, or deep-learning approaches – such as graph neural networks – to further improve precision and robustness in future implementations. In addition, the isotope/adduct detection algorithm and its underlying concept could be further developed by incorporating additional chemical constraints, improved mass-difference modeling, more advanced learning-based classification methods, and real-time integration with online annotation platforms such as METASPACE [66] to enhance the accuracy of peak relationship identification in MALDI-MSI datasets.

## Acknowledgements

The mouse strain used for this research project, B6;129-Tg(APPSwe,tauP301L)1Lfa *Psen1*^*tm1Mpm*^/Mmjax, RRID:MMRRC_034830-JAX, was obtained from the Mutant Mouse Resource and Research Center (MMRRC) at The Jackson Laboratory, an NIH-funded strain repository, and was donated to the MMRRC by Frank Laferla, Ph.D., University of California, Irvine, and Mark P. Mattson, Ph.D., Johns Hopkins University, School of Medicine.

Funding for animals was provided by UT-Oak Ridge Innovation Institute (UT-ORII) Science Alliance: Support for Affiliated Research Teams (StART) Program. Amin Jarrahi was supported by UTK ORIED Human Health and Wellness Gateway FY2026 and the ORIED AI Tennessee Initiative Seed Funds.

## Footnotes

a https://www.biorender.com

b Some parts of the flowchart was adopted from https://www.bmkgene.com/10x-genomics-visium-spatial-transcriptome-product/

c The image is produced by AI (https://chatgpt.com)

d https://www.biorender.com

## Notes

### Competing Interest Statement

The authors have declared no competing interest.

## References

[1] M. Wess et al., “Spatial integration of multi-omics data from serial sections using the novel Multi-Omics Imaging Integration Toolset,” Gigascience, vol. 14, pp. 1–19, Jan. 2025, doi: 10.1093/gigascience/giaf035.

[2] A. Enninful et al., “Integration of imaging-based and sequencing-based spatial omics mapping on the same tissue section via DBiTplus,” Nature Methods 2026, pp. 1–13, Jan. 2026, doi: 10.1038/s41592-025-02948-0.

[3] T. F. E. Hendriks, G. B. Eijkel, T. Visvikis, B. Balluff, R. M. A. Heeren, and E. Cuypers, “One section, two worlds: single-cell integration of MALDI-MSI and spatial transcriptomics on the same single tissue section,” Scientific Reports 2025 15:1, vol. 15, no. 1, pp. 42660-, Nov. 2025, doi: 10.1038/s41598-025-26735-1.

[4] E. C. Williams et al., “Spatially resolved integrative analysis of transcriptomic and metabolomic changes in tissue injury studies,” Nature Communications 2026 17:1, vol. 17, no. 1, pp. 205-, Jan. 2026, doi: 10.1038/s41467-025-68003-w.

[5] X. Liu et al., “Spatial multi-omics: deciphering technological landscape of integration of multiomics and its applications,” Journal of Hematology & Oncology 2024 17:1, vol. 17, no. 1, pp. 72-, Aug. 2024, doi: 10.1186/s13045-024-01596-9.

[6] J. F. Navarro et al., “Spatial Transcriptomics Reveals Genes Associated with Dysregulated Mitochondrial Functions and Stress Signaling in Alzheimer Disease,” iScience, vol. 23, no. 10, Oct. 2020, doi: 10.1016/j.isci.2020.101556.

[7] T. C. Gripshover et al., “Visium spatial transcriptomics and proteomics identifies novel hepatic cell populations and transcriptomic signatures of alcohol-associated hepatitis,” Alcohol: Clinical and Experimental Research, vol. 49, no. 1, pp. 106–116, Jan. 2025, doi: 10.1111/acer.15494.

[8] B. Heijs, E. A. Tolner, J.V. M. G. Bovée, A. M. J. M. Van Den Maagdenberg, and L. A. McDonnell, “Brain Region-Specific Dynamics of On-Tissue Protein Digestion Using MALDI Mass Spectrometry Imaging,” J. Proteome Res., vol. 14, no. 12, pp. 5348–5354, Dec. 2015, doi: 10.1021/acs.jproteome.5b00849.

[9] R. T. Steven and J. Bunch, “Repeat MALDI MS imaging of a single tissue section using multiple matrices and tissue washes,” Analytical and Bioanalytical Chemistry 2013 405:14, vol. 405, no. 14, pp. 4719–4728, Mar. 2013, doi: 10.1007/s00216-013-6899-9.

[10] M. Stoeckli, D. Staab, M. Staufenbiel, K. H. Wiederhold, and L. Signor, “Molecular imaging of amyloid β peptides in mouse brain sections using mass spectrometry,” Anal. Biochem., vol. 311, no. 1, pp. 33–39, Dec. 2002, doi: 10.1016/S0003-2697(02)00386-X.

[11] S. Ntshangase, S. Mdanda, T. Naicker, H. G. Kruger, S. Baijnath, and T. Govender, “Spatial distribution of elvitegravir and tenofovir in rat brain tissue: Application of matrix-assisted laser desorption/ionization mass spectrometry imaging and liquid chromatography/tandem mass spectrometry,” Rapid Communications in Mass Spectrometry, vol. 33, no. 21, pp. 1643–1651, Nov. 2019, doi: 10.1002/rcm.8510.

[12] Y. Miyake, S. Kusaka, I. Murata, and M. Toyoda, “Matrix-Assisted Laser Desorption/Ionization (MALDI) Mass Spectrometry Imaging of L-4-Phenylalanineboronic Acid (BPA) in a Brain Tumor Model Rat for Boron Neutron Capture Therapy (BNCT),” Mass Spectrometry, vol. 11, no. 1, pp. A0105–A0105, Dec. 2022, doi: 10.5702/massspectrometry.A0105.

[13] C. D. Cerruti, F. Benabdellah, O. Laprévote, D. Touboul, and A. Brunelle, “MALDI Imaging and Structural Analysis of Rat Brain Lipid Negative Ions with 9-Aminoacridine Matrix,” Anal. Chem., vol. 84, no. 5, pp. 2164–2171, Mar. 2012, doi: 10.1021/ac2025317.

[14] R. Argelaguet et al., “MultiCOmics Factor Analysis—a framework for unsupervised integration of multiComics data sets,” Molecular Systems Biology 2018 14:6, vol. 14, no. 6, pp. MSB178124-, Jun. 2018, doi: 10.15252/msb.20178124.

[15] R. Argelaguet et al., “MOFA+: a statistical framework for comprehensive integration of multi-modal single-cell data,” Genome Biology 2020 21:1, vol. 21, no. 1, pp. 111-, May 2020, doi: 10.1186/s13059-020-02015-1.

[16] Y. Zhou, X. Xiao, L. Dong, C. Tang, G. Xiao, and L. Xu, “Cooperative integration of spatially resolved multi-omics data with COSMOS,” Nature Communications 2024 16:1, vol. 16, no. 1, pp. 27-, Jan. 2025, doi: 10.1038/s41467-024-55204-y.

[17] R. M. Mersereau, “The Processing of Hexagonally Sampled Two-Dimensional Signals,” Proceedings of the IEEE, vol. 67, no. 6, pp. 930–949, 1979, doi: 10.1109/PROC.1979.11356.

[18] W. E. Snyder, H. Qi, and W. A. Sander, “Coordinate system for hexagonal pixels,” in Proceeding Volume 3661, Medical Imaging, Image Processing, San Diego, CA: Society of Photo-Optical Instrumentation Engineers (SPIE), May 1999, pp. 716–727. doi: 10.1117/12.348629.

[19] J. A. Soria Lopez, H. M. González, and G. C. Léger, “Alzheimer’s disease,” Handb. Clin. Neurol., vol. 167, pp. 231–255, Jan. 2019, doi: 10.1016/B978-0-12-804766-8.00013-3.

[20] M. W. Bondi, E. C. Edmonds, and D. P. Salmon, “Alzheimer’s Disease: Past, Present, and Future,” Journal of the International Neuropsychological Society, vol. 23, no. 9–10, pp. 818– 831, Oct. 2017, doi: 10.1017/S135561771700100X.

[21] I. Kaya et al., “Novel Trimodal MALDI Imaging Mass Spectrometry (IMS3) at 10 μm Reveals Spatial Lipid and Peptide Correlates Implicated in Aβ Plaque Pathology in Alzheimer’s Disease,” ACS Chem. Neurosci., vol. 8, no. 12, pp. 2778–2790, Dec. 2017, doi: 10.1021/acschemneuro.7b00314.

[22] N. Kakuda et al., “Distinct deposition of amyloid-β species in brains with Alzheimer’s disease pathology visualized with MALDI imaging mass spectrometry,” Acta Neuropathologica Communications 2017 5:1, vol. 5, no. 1, pp. 73-, Oct. 2017, doi: 10.1186/s40478-017-0477-x.

[23] S. Oddo et al., “Triple-transgenic model of Alzheimer’s Disease with plaques and tangles: Intracellular Aβ and synaptic dysfunction,” Neuron, vol. 39, no. 3, pp. 409–421, Jul. 2003, doi: 10.1016/S0896-6273(03)00434-3.

[24] A. T. Du et al., “Magnetic resonance imaging of the entorhinal cortex and hippocampus in mild cognitive impairment and Alzheimer’s disease,” J. Neurol. Neurosurg. Psychiatry, vol. 71, no. 4, pp. 441–447, Oct. 2001, doi: 10.1136/jnnp.71.4.441.

[25] D. Wang et al., “Elevated plasma levels of exosomal BACE1-AS combined with the volume and thickness of the right entorhinal cortex may serve as a biomarker for the detection of Alzheimer’s disease,” Mol. Med. Rep., vol. 22, no. 1, pp. 227–238, Jul. 2020, doi: 10.3892/mmr.2020.11118.

[26] E. Schueller et al., “Dysregulation of histone acetylation pathways in hippocampus and frontal cortex of Alzheimer’s disease patients,” European Neuropsychopharmacology, vol. 33, pp. 101–116, Apr. 2020, doi: 10.1016/j.euroneuro.2020.01.015.

[27] M. Dufresne, L. G. Migas, K. V. Djambazova, M. E. Colley, R. Van de Plas, and J. M. Spraggins, “Aminated cinnamic acid analogs as dual polarity matrices for high spatial resolution MALDI imaging mass spectrometry,” Anal. Chim. Acta, vol. 1371, no. 23, p. 344423, Oct. 2025, doi: 10.1016/j.aca.2025.344423.

[28] A. Jarrahi, A. Jones, W. Tang, H. Qi, and A. C. Crouch, “Mathematical Framework for Quantifying Delocalization in MALDI-MSI via a Composite Scoring Approach,” ACS Measurement Science Au, vol. 6, p. 58, Jan. 2026, doi: 10.1021/acsmeasuresciau.5c00148.

[29] A. Savitzky and M. J. E. Golay, “Smoothing and Differentiation of Data by Simplified Least Squares Procedures,” Anal. Chem., vol. 36, no. 8, pp. 1627–1639, Jul. 1964, doi: 10.1021/AC60214A047/ASSET/AC60214A047.FP.PNG_V03.

[30] P. H. C. Eilers and H. F. M. Boelens, “Baseline Correction with Asymmetric Least Squares Smoothing,” Leiden University Medical Centre Report, vol. 1, no. 1, p. 5, Oct. 2005.

[31] D. C. Castro et al., “Single-Cell and Subcellular Analysis Using Ultrahigh Resolution 21 T MALDI FTICR Mass Spectrometry,” Anal. Chem., vol. 95, no. 17, pp. 6980–6988, May 2023, doi: 10.1021/acs.analchem.3c00393.

[32] M. Blank, T. Enzlein, and C. Hopf, “LPS-induced lipid alterations in microglia revealed by MALDI mass spectrometry-based cell fingerprinting in neuroinflammation studies,” Scientific Reports 2022 12:1, vol. 12, no. 1, pp. 2908-, Feb. 2022, doi: 10.1038/s41598-022-06894-1.

[33] Z. Hall et al., “MYC expression drives aberrant lipid metabolism in lung cancer,” Cancer Res., vol. 76, no. 16, pp. 4608–4618, Aug. 2016, doi: 10.1158/0008-5472.CAN-15-3403.

[34] Y. Otsuka et al., “Improved ion detection sensitivity in mass spectrometry imaging using tapping-mode scanning probe electrospray ionization to visualize localized lipids in mouse testes,” Analytical and Bioanalytical Chemistry 2024 417:2, vol. 417, no. 2, pp. 275–286, Nov. 2024, doi: 10.1007/s00216-024-05641-x.

[35] A. Shenfeld and A. Galkin, “Role of the MBP protein in myelin formation and degradation in the brain,” Biological Communications, vol. 67, no. 2, pp. 127–138, Jun. 2022, doi: 10.21638/spbu03.2022.206.

[36] J. H. Ahn et al., “Age-dependent differences in myelin basic protein expression in the hippocampus of young, adult and aged gerbils,” Lab. Anim. Res., vol. 33, no. 3, p. 237, Jul. 2017, doi: 10.5625/lar.2017.33.3.237.

[37] C. Dimovasili et al., “Aging compromises oligodendrocyte precursor cell maturation and efficient remyelination in the monkey brain,” Geroscience, vol. 45, no. 1, p. 249, Feb. 2022, doi: 10.1007/s11357-022-00621-4.

[38] M. Gadek et al., “Aging activates escape of the silent X chromosome in the female mouse hippocampus,” Science Advances, vol. 11, no. 10, p. 8169, Mar. 2025, doi: 10.1126/sciadv.ads8169.

[39] J. Wang et al., “S100B gene polymorphisms are associated with the S100B level and Alzheimer’s disease risk by altering the miRNA binding capacity,” Aging, vol. 13, no. 10, pp. 13954–13967, May 2021, doi: 10.18632/aging.203005.

[40] N. V. Bobkova et al., “Exogenous Hsp70 delays senescence and improves cognitive function in aging mice,” Proc. Natl. Acad. Sci. U. S. A., vol. 112, no. 52, pp. 16006–16011, Dec. 2015, doi: 10.1073/pnas.1516131112.

[41] M. Ximerakis et al., “Heterochronic parabiosis reprograms the mouse brain transcriptome by shifting aging signatures in multiple cell types,” Nature Aging 2023 3:3, vol. 3, no. 3, pp. 327– 345, Mar. 2023, doi: 10.1038/s43587-023-00373-6.

[42] M. Babić Leko et al., “Further validation of the association between MAPT haplotype-tagging polymorphisms and Alzheimer’s disease: neuropsychological tests, cerebrospinal fluid biomarkers, and APOE genotype,” Front. Mol. Neurosci., vol. 17, p. 1456670, Sep. 2024, doi: 10.3389/fnmol.2024.1456670.

[43] C. C. Zhang, A. Xing, M. S. Tan, L. Tan, and J. T. Yu, “The Role of MAPT in Neurodegenerative Diseases: Genetics, Mechanisms and Therapy,” Molecular Neurobiology 2015 53:7, vol. 53, no. 7, pp. 4893–4904, Sep. 2015, doi: 10.1007/s12035-015-9415-8.

[44] H. Mori et al., “Blood MAPT expression and methylation status in Alzheimer’s disease,” Psychiatry and Clinical Neurosciences Reports, vol. 1, no. 4, p. e65, Dec. 2022, doi: 10.1002/pcn5.65.

[45] K. Schindowski et al., “Alzheimer’s Disease-Like Tau Neuropathology Leads to Memory Deficits and Loss of Functional Synapses in a Novel Mutated Tau Transgenic Mouse without Any Motor Deficits,” Am. J. Pathol., vol. 169, no. 2, p. 599, 2006, doi: 10.2353/ajpath.2006.060002.

[46] N. Bin Abid, M. I. Naseer, and M. O. Kim, “Comparative Gene-Expression Analysis of Alzheimer’s Disease Progression with Aging in Transgenic Mouse Model,” Int. J. Mol. Sci., vol. 20, no. 5, p. 1219, Mar. 2019, doi: 10.3390/ijms20051219.

[47] C. Bellenguez et al., “New insights into the genetic etiology of Alzheimer’s disease and related dementias,” Nature Genetics 2022 54:4, vol. 54, no. 4, pp. 412–436, Apr. 2022, doi: 10.1038/s41588-022-01024-z.

[48] E. Bellou et al., “Age-dependent effect of APOE and polygenic component on Alzheimer’s disease,” Neurobiol. Aging, vol. 93, p. 69, Sep. 2020, doi: 10.1016/j.neurobiolaging.2020.04.024.

[49] A. Labuza et al., “Aging, Rather than Genotype, Is the Principal Contributor to Differential Gene Expression Within Targeted Replacement APOE2, APOE3, and APOE4 Mouse Brain,” Brain Sci., vol. 15, no. 10, p. 1117, Oct. 2025, doi: 10.3390/brainsci15101117.

[50] A. C. Raulin, S. V. Doss, Z. A. Trottier, T. C. Ikezu, G. Bu, and C. C. Liu, “ApoE in Alzheimer’s disease: pathophysiology and therapeutic strategies,” Molecular Neurodegeneration 2022 17:1, vol. 17, no. 1, pp. 72-, Nov. 2022, doi: 10.1186/s13024-022-00574-4.

[51] O. Catz and M. B. Lewis, “Exploring distinctiveness, attractiveness and sexual dimorphism in actualized face-spaces,” Vis. cogn., vol. 28, no. 9, pp. 453–469, Oct. 2020, doi: 10.1080/13506285.2020.1797967.

[52] T. Valentine, “A Unified Account of the Effects of Distinctiveness, Inversion, and Race in Face Recognition,” The Quarterly Journal of Experimental Psychology Section A, vol. 43, no. 2, pp. 161–204, May 1991, doi: 10.1080/14640749108400966.

[53] K. Pearson, “VII. Note on regression and inheritance in the case of two parents,” Proceedings of the Royal Society of London, vol. 58, no. 347–352, pp. 240–242, Dec. 1895, doi: 10.1098/rspl.1895.0041.

[54] P. A. P. Moran, “Notes on Continuous Stochastic Phenomena,” Biometrika, vol. 37, no. 1/2, p. 17, Jun. 1950, doi: 10.2307/2332142.

[55] L. Anselin, “Local Indicators of Spatial Association—LISA,” Geogr. Anal., vol. 27, no. 2, pp. 93–115, Apr. 1995, doi: 10.1111/j.1538-4632.1995.tb00338.x.

[56] M. J. D. Powell, “An efficient method for finding the minimum of a function of several variables without calculating derivatives,” Comput. J., vol. 7, no. 2, pp. 155–162, Jan. 1964, doi: 10.1093/comjnl/7.2.155.

[57] P. Virtanen et al., “SciPy 1.0: fundamental algorithms for scientific computing in Python,” Nature Methods 2020 17:3, vol. 17, no. 3, pp. 261–272, Feb. 2020, doi: 10.1038/s41592-019-0686-2.

[58] M. Höring et al., “Correction of Isobaric Overlap Resulting from Sodiated Ions in Lipidomics,” Anal. Chem., vol. 92, no. 16, pp. 10966–10970, Aug. 2020, doi: 10.1021/acs.analchem.0c02408.

[59] J. Dehairs et al., “LipidQMap - An Open-Source Tool for Quantitative Mass Spectrometry Imaging of Lipids,” bioRxiv, p. 2025.10.15.682422, Oct. 2025, doi: 10.1101/2025.10.15.682422.

[60] Y. Miyamoto, T. Torii, A. Tanoue, and J. Yamauchi, “Pelizaeus–Merzbacher disease-associated proteolipid protein 1 inhibits oligodendrocyte precursor cell differentiation via extracellular-signal regulated kinase signaling,” Biochem. Biophys. Res. Commun., vol. 424, no. 2, pp. 262–268, Jul. 2012, doi: 10.1016/j.bbrc.2012.06.101.

[61] D. L. Fulton, E. Denarier, H. C. Friedman, W. W. Wasserman, and A. C. Peterson, “Towards resolving the transcription factor network controlling myelin gene expression,” Nucleic Acids Res., vol. 39, no. 18, pp. 7974–7991, Oct. 2011, doi: 10.1093/nar/gkr326.

[62] F. Michetti et al., “The S100B story: from biomarker to active factor in neural injury,” J. Neurochem., vol. 148, no. 2, pp. 168–187, Jan. 2019, doi: 10.1111/jnc.14574.

[63] J. Du et al., “S100B is selectively expressed by gray matter protoplasmic astrocytes and myelinating oligodendrocytes in the developing CNS,” Mol. Brain, vol. 14, no. 1, p. 154, Dec. 2021, doi: 10.1186/s13041-021-00865-9.

[64] J. Steiner et al., “Evidence for a wide extra-astrocytic distribution of S100B in human brain,” BMC Neuroscience 2007 8:1, vol. 8, no. 1, pp. 2-, Jan. 2007, doi: 10.1186/1471-2202-8-2.

[65] G. J. Siegel, R. Wayne Albers, S. T. Brady, and D. L. Price, Eds., Basic Neurochemistry: Molecular, Cellular and Medical Aspects, 7th ed. Amsterdam, The Netherlands: Elsevier Academic Press, 2006.

[66] D. D. Nguyen et al., “Facilitating imaging mass spectrometry of microbial specialized metabolites with METASPACE,” Metabolites, vol. 11, no. 8, p. 477, Aug. 2021, doi: 10.3390/metabo11080477.

